# Functional details of the integral membrane metallo-protease FtsH revealed by solution NMR spectroscopy

**DOI:** 10.64898/2026.05.06.723347

**Authors:** Hannah Fremlén, Björn M. Burmann

**Affiliations:** Department of Chemistry and Molecular Biology, University of Gothenburg, 405 30 Göteborg, Sweden; Wallenberg Centre for Molecular and Translational Medicine, University of Gothenburg, 405 30 Göteborg, Sweden; Swedish NMR Centre, University of Gothenburg, 405 30 Göteborg, Sweden; Science for Life Laboratory, Integrated Structural Biology Platform, University of Gothenburg, 405 30 Göteborg, Sweden

**Author notes:** Correspondence should be addressed to BMB.

## Abstract

The bacterial FtsH protease resides in the inner leaflet of the bacterial inner membrane and is the sole essential protease found in *Escherichia Coli* (*E. coli*). FtsH’s mandatory role is due to its proteolytic regulation of a key enzyme in the lipopolysaccharide (LPS) assembly, LpxC, although FtsH substrates comprise a diverse pool of both soluble and membrane proteins. While the full-length structure of FtsH could be solved recently, large parts of its functional cycle remain poorly understood. Here, we use advanced solution NMR spectroscopy methods in combination with biochemical assays to elucidate its ATPase cycle at an unprecedented level of detail. We reveal the foundation of the interdomain communication based on subtly tuned protein dynamics mediated *via* key NOE contacts as well as the kinetic rates underlying ATP consumption, providing a detailed level of insight into the interplay between the individual FtsH sub-domains, that drive the proteolytic cycle.

## Introduction

Cells in all organisms depend on regulated proteolysis to maintain a healthy proteome, as the accumulation of misfolded and aggregated proteins potentially can be lethal (*1*, *2*). Cellular protein quality control relies heavily on a family of AAA+ proteases (ATPases associated with various cellular activities) to carry out the task of homeostasis (*3*, *4*), and maintaining a functional proteome. AAA+ proteases are large, self-compartmentalizing machineries forming ring-like structures and using the energy released during ATP-consumption to drive unfolding and subsequent translocation of its substrates for degradation (*4*). *Escherichia coli* (*E. coli*) harbors the most well-studied AAA+ proteases of which the integral membrane protein FtsH (filamentous temperature-sensitive H) is the only essential one for cell survival (*5*, *6*). Localized to the inner bacterial membrane, FtsH degrades both membrane-bound as well as cytosolic proteins (*7*). The essential role of FtsH stems from its involvement in the regulation of the import of lipopolysaccharides (LPS) and phospholipids (PL) to the outer leaflet of the outer membrane by degradation of LpxC, a key enzyme in the LPS synthesis pathway (*8*).

Structurally, FtsH assembles into a ring-like homohexamer harboring smaller periplasmic domains, in total 12 membrane-spanning α-helices and a larger cytoplasmic domain carrying its essential functionality (*9*). Both the AAA+ domain and the protease domain are found on a single polypeptide chain and make up the cytoplasmic domain (*10*). The AAA+ domain, responsible for unfolding and translocation of substrates, encompasses several conserved, characteristic motifs found in the AAA+ family of proteases. These include the Walker A- and Walker B box, involved in nucleotide binding and hydrolysis, and the second region of homology (SRH) encompassing “the arginine finger”, crucial for ATP hydrolysis (*11*). The protease domain carries a zinc-binding motif, HEXXH (X refers to any residue) characteristic for metalloproteases in which the histidines together with an adjacent glutamic acid coordinate Zn^2+^ ions required for proteolytic activity (*10*, *12*).

Homologs to FtsH can be found across all domains of life, and serve important roles in for example mitochondrial protein quality control, and mutations in the human m-AAA protease are associated with neurodegenerative diseases (*8*, *13*). Previous structural studies of FtsH and its homologs have mainly been focused on isolated, soluble domains. Both the AAA+ domain and the protease domain display mainly an α-helical fold, with small segments of β-strands. While the protease domain forms a symmetric hexameric-ring structure, the AAA+ domain seems to be more dynamic and is formed by two subdomains (*11*). More recent studies conducted by cryo-electron microscopy (cryo-EM) show a large complex of the full-length protein together with another two other membrane proteins, HflK and HflC at medium resolution (*14*). HflKC is thought to act as a regulator, inhibiting the activity of FtsH based on cellular environment.

Here we set out to specify the functional details of the FtsH ATPase cycle. Using a combination of solution NMR spectroscopy approaches, proteolytic cleavage and an array of functional ATPase assay, we reveal the steps of the ATP cycle at unprecedented detail and elucidate the functional interplay between ATPase and protease domain within FtsH. In order to gain more insight into the mechanistic details, we adopted an approach recently used for a homologous mitochondrial protein, YME1L, employing a construct of the functional domains fused to a peptide which forms a hexameric coiled coil in solution, ccHex (*15*). Thus, by replacing the transmembrane segment and obtaining a soluble hexameric, active construct, we were able to study FtsH function in unprecedented detail. Using a combination of protein dynamics, methyl NOE patterns, *in-cyclo* NMR alongside classical ATP-functional assays, enabled us to derive details of allosteric pathways highlighting the control of the ATPase function by the proteolytic site by limiting the speed of the ATP hydrolysis.

## Methods

### Expression and Purification

A construct of the soluble cytoplasmic domain of FtsH (138–644), lacking the transmembrane and periplasmic segments, (cFtsH) in a pET28b(+) vector with an amino-terminal His_6_-tag was purchased from GenScript. Using restriction-free cloning (*16*), the AAA+ domain (139–398, aFtsH) and the protease domain (399–644, pFtsH) residing within the cytoplasmic domain, were cloned separately into a pET28b(+), which also included an N-terminal SUMO-tag following the His_6_-tag. Finally, cFtsH was cloned into the ccHex vector obtained from GenScript (SUMO-ccHex in a pet28b(+)-vector) resulting in the hexFtsH construct using restriction-free cloning. Point mutations to create constructs with impaired ATPase activity (hexFtsH_K201N_) and proteolytic activity (hexFtsH_E415Q_) were also performed *via* restriction-free cloning. All used primer sequences and used plasmids are provided in **tables S1** and **S2**.

All constructs were expressed in *E. coli* BL21 (λDE3) cells. Unlabeled FtsH samples were expressed in LB media, while isotope labeled samples were grown in minimal media (M9), both supplemented with 50 μg/ml kanamycin. For all constructs, except for aFtsH, the M9 medium was also enriched with 100 μM ZnSO_4_. Cells were grown at 37°C and induced with 0.5 mM IPTG (isopropyl β-D-1-thiogalactopyranoside) at an OD_600_ of ∼0.8. Protein expression was continued at 20°C for an additional 12–16 hours. The cell cultures were harvested by centrifugation at 4.000 x g for 30 minutes at 4°C. Thereafter, the cells were resuspended in the respective lysis buffer, where aFtsH, pFtsH, cFtsH were resuspended in buffer A (20 mM Tris, 500 mM NaCl, 1 mM DTT and 5 mM Imidazole, pH 8) and hexFtsH in buffer B (50 mM Tris, 300 mM NaCl, 1 mM DTT, 10% glycerol, pH 8). Prior to lysis, a protease inhibitor cocktail tablet (Roche), ∼125 U DNase (ArcticZymes) and 5 mM MgCl_2_ were added to the sample. Cells were lysed using a high-pressure homogenizer (Emulsiflex, Avestin) and subsequently centrifuged at 18.000 x g for 1 hour at 4°C before being loaded onto a Ni^2+^-HisTrap column (GE Healthcare) pre-equilibrated with the respective lysis buffer. Elution from the column was conducted with 500 mM imidazole added to the lysis buffer. Fractions containing aFtsH, pFtsH and hexFtsH were pooled and dialyzed overnight at 4°C against respective lysis buffer to cleave the SUMO-tag upon addition of Senp1 (Addgene #16356) (*17*). The samples were subsequently re-applied to a Ni^2+^-HisTrap column to elute the now tagless samples. aFtsH, pFtsH, cFtsH and hexFtsH were in a next step concentrated using Vivaspin concentrators (10K MWCO for aFtsH and pFtsH, respectively, 30K MWCO for cFtsH, and 50K MWCO for hexFtsH FF; Sartorius) and then applied to a Superdex 200 Increase column (GE Healthcare) equilibrated with buffer C (50 mM potassium phosphate, 300 mM KCl, pH 7.4) or buffer D (20 mM Tris, 150 mM NaCl, 5 mM MgCl_2_, 1 mM DTT, 10% glycerol, pH 7.8) for hexFtsH. Purity was assessed by SDS PAGE using 4–20% MiniProtean TGX gels (Bio-Rad) and the purified proteins were stored at -80°C until usage. All buffer compositions used are provide in **table S3.**

After the size-exclusion step, hexFtsH was incubated with 2 units (U) of apyrase (New England Biolabs) and 5 mM CaCl_2_, in buffer D, overnight at room temperature (∼21 °C) to ensure complete digestion of any remaining endogenously bound nucleotides. The apyrase treated samples were then re-purified on a Superdex 200 Increase column (GE Healthcare) equilibrated with buffer D to remove apyrase from hexFtsH. Protein containing fractions were pooled, flash frozen in liquid nitrogen, and stored at -80°C until usage.

### Isotope Labelling

NMR samples were isotope labeled by addition of relevant isotopes to the M9 media during expression. H_2_O-based media were supplemented either with (^15^NH_4_)Cl for [*U*-^15^N]-labeled protein or with both (^15^NH_4_)Cl and *D*-(^13^C)-glucose for uniform double labeling for [*U*-^13^C,^15^N]. For proteins expressed in D_2_O-based M9 media addition of (^15^NH_4_)Cl yielded [*U*-^2^H,^15^N]-labelling whereas addition of (^15^NH_4_)Cl and *D*-(^2^H,^13^C)-glucose resulted in [*U*-^2^H,^13^C,^15^N]-labelled proteins.

For specific methyl labeled samples of cFtsH and hexFtsH, the D_2_O-based M9 medium was supplemented with (^15^NH_4_)Cl and *D*-(^2^H,^12^C)-glucose together with the respective amino acid or metabolic precursor. The precursors were added to the culture 1 h prior to induction. For the methionine and isoleucine labeled sample, 50 mg/ml [*U*-^2^H,^13^C,^13^] methionine alongside 50 mg/ml 2-Ketobutyric acid-4-^13^C,3,3-d_2_ sodium salt hydrate (isoleucine) was added. The alanine sample was labeled using 50 mg/l 2-[^2^H], 3-[^13^C] L-alanine and 2% Bioexpress to suppress isotope scrambling (*18*). For *proS* stereo-specific isotope labelling of valine/leucine, the DLAM-LV*^proS^*-kit (2-(^13^C)-methyl-4-(D_3_)-acetolactate was added. Bioexpress and alanine were purchased from Cambridge Isotopes Laboratories, DLAM-LV*^proS^* precursors from NMR-Bio. All other isotopes were purchased from Merck.

### NMR-Spectroscopy

NMR experiments were recorded at 310 K on Bruker Avance III HD 700, 800 or 900 MHz spectrometers equipped with cryogenic cooled triple- or quadruple-resonance probes, running Topspin 3.6 (Bruker Biospin). Backbone resonance assignment of aFtsH and pFtsH were obtained using the following TROSY-type 3D experiments; HNCO, HNCACO, HNCOCA, HNCA, HNCACB and CBCACONH (*19*, *20*). Aliphatic side-chain resonance assignments of the same constructs were achieved by 3D (H)C(CCO)NH, H(CCCO)NH as well as HBHACONH, and based on 2D [^13^C,^1^H]-HSQC spectra with/without constant time (CT) evolution by 3D HC(C)H-TOCSY and 3D (H)CCH-TOCSY experiments (*21*). Furthermore, assignment of methyl groups of cFtsH was facilitated by 3D ^13^C_methyl_-^13^C_methyl_-^1^H_methyl_ SOFAST NOESY and 3D ^1^H_methyl_-^15^N_amide_-^1^H_amide_ SOFAST NOESY experiments using 300 ms mixing time in both cases (*22*). Processing of NMR data were performed using a combination of the mddNMR (*23*) and NMRPipe (*24*) software packages. Sequence specific resonance assignments and spectral analyses were done in CARA (*25*). The secondary chemical shifts of the backbone were calculated relative to the random coil values derived by the POTENCI algorithm (*26*). For the obtained raw data, a weighting function with weights *1-2-1* for residues *(i-1)-i-(i+1)* was applied (*27*, *28*).

The chemical shift changes of the side-chain carbons were calculated according to Equation 1.

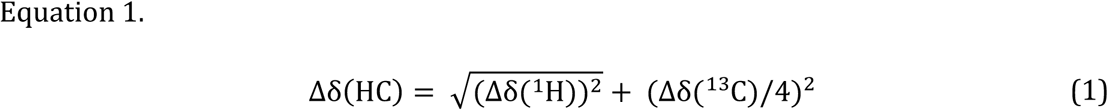

For quantitative analysis of signal intensities, the signal amplitudes were corrected by dilution factor, number of scans and differences in the ^1^H-90° pulse length (*29*).

We used the obtained Ile (δ_1_), and Met (ε) ^13^C chemical shifts to deduce the rotameric equilibria of the Ile ξ_2_ and Met ξ_3_ angles employing a previously outlined approach (*30*, *31*). The ^13^C chemical shifts of methyl groups are directly dependent on the side chain rotamer, thus this approach yields direct insight into the rotameric state of the different methyl groups (*32*, *33*). The population of the *trans* rotameric state (*p*_trans_) for each residue was calculated according to the chemical shift values of the methyl ^13^C signals (*δ*_obs_: ppm) using equations (2 and 3):

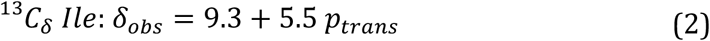

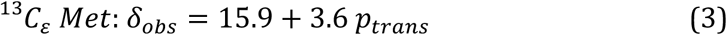

*p*_trans_ values determined by these equations ranged from 1 (all *trans*) to 0 (all *gauche –*), respectively. Residues with values ranging between 0.75 – 1 were considered to be in the *trans* conformation, whereas those between 0 – 0.25 were in the *gauche –* conformation.

### NMR backbone dynamics

Backbone relaxation data for aFtsH and pFtsH were obtained through ^15^N{^1^H}-NOE, *R*_1_(^15^N) and *R*_1π_(^15^N) (*34*), recorded at 800 MHz (18.8 T) as well as 900 MHz (21.1 T). Non-linear least square fits of the relaxation data were done using PINT (*35*). *R*_2(R1π)_ ^15^N values were extracted from the obtained *R* _1ρ_ rates using Equation 4.

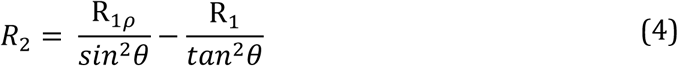

with θ = tan^-1^(ω/Ω), where ω is the spin-lock field strength (2 kHz) and Ω is the offset from the ^15^N carrier frequency.

Error bars for *^1^*^5^N{^1^H}-NOE were calculated from the spectral noise and error bars for *R*_1_(^15^N) and *R*_1ρ_(^15^N) were calculated by a Monte Carlo simulation embedded in PINT (*35*) whereas the *R*_2(R1ρ)_ (^15^N) errors were calculated by error propagation. Analysis of the resulting relaxation rates was performed using Tensor2 (*36*) hosted on the NMRbox web server (*37*). The relaxation rates were further analysed using FASTModelfree (*38*) with the aFtsH crystal structure (PDB-ID: 1LV7) and the AlphaFold3 model for pFtsH (AlphaFoldDB: P0AAI3) as structural templates. Theoretical relaxation rates were obtained from HYDRONMR version 7c (*39*) *via* NMRBox using a standard atomic element radius value of 3.3, a 1.02 Å N–H bond length and a Chemical Shift Anisotropy (CSA) of 172 ppm. Subsequently we performed further analysis of the obtained relaxation rates using the pyDIFRATE program, a python-based tool for detectors analysis (https://github.com/alsinmr/pyDR) (*40*, *41*). Three to four detectors were used for the analysis, which provided similar results.

### NMR sidechain dynamics

Experiments were performed on separate cFtsH-samples with the following labelling schemes; [*U*-^2^H, Ala-^13^CH_3_], [*U*-^2^H, Ile-δ_1_-^13^CH_3_, Met-ε-^13^CH_3_] and [*U*-^2^H, Leu,Val*^proS^*-^13^CH_3_] (*42*), at a temperature of 310 K in 99.9% D_2_O based NMR buffer. Sidechain methyl order parameters (S^2^_axis_•τ_c_) were determined by cross-correlated relaxation experiments (*43*, *44*). Single- (SQ) and triple-quantum (TQ) ^1^H-^13^C experiments were collected at a series of delays. Ratios of peak intensities were determined in PINT (*35*) and subsequently fitted for 19 relaxation delay times ranging between 1.1 ms to 32 ms using Equation 5:

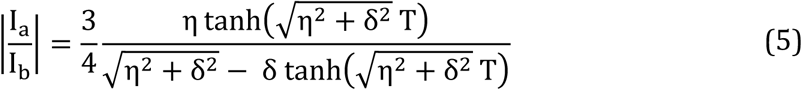

where *T* is the relaxation delay time and δ is a factor to account for coupling due to relaxation with external protons.

S^2^_axis_•τ_c_ values were determined using Equation 6, using their fitted η values.

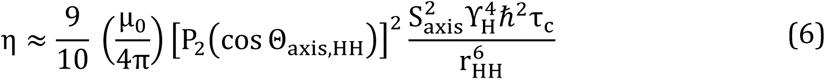

where μ_0_ is the vacuum permittivity constant, γ_H_ is the gyromagnetic ratio of the proton spin, *r*_HH_ is the distance between pairs of methyl protons (1.813 Å), S^2^_axis_ is the generalized order parameter describing the amplitude of motion of the methyl threefold axis, Θ_axis,HH_ is the angle between the methyl symmetry axis and a vector between pair of methyl protons (90°), and P_2_(x) = ½(3x^2^-1). Data were analysed by in-house written scripts and the product of the methyl order parameter and the overall correlation time constant, S^2^_axis_•τ_c_, was determined.

Multiple quantum (MQ) methyl relaxation dispersion experiments were recorded as a series of 2D datasets using constant time relaxation periods (*T*) of 40 ms (16.4 T (700 MHz)/ 21.1 T (900 MHz)) and CPMG (Carr-Purcell-Meiboom-Gill) frequencies ranging from 33 to 1.000 Hz (*45*). The effective transverse relaxation rate, *R*_2,eff_, was calculated according to the following equation:

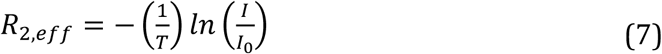

where *I* (*I*_0_) are the intensities with (without) the presence of a constant-time relaxation interval of duration *T*, during which a variable number of ^13^C 180° pulses are applied leading to ν_CPMG_ = 1/(2δ), where δ is the time between successive pulses. Data were processed in nmrPipe (*24*), and the peak intensities were extracted with PINT (*35*).

### In-Gel proteolytic cleavage assay

All cleavage assays were performed at 37°C with 0.6 μM FtsH (hexamer concentration) in 25 mM HEPES, 150 mM KCl, 5 mM MgCl_2_, 25 μM ZnCl_2_, 1 mM DTT, pH 7.4 (Buffer E) supplemented with either 2 mM ATP (Sigma-Aldrich) or 2 mM ATPγS (Sigma-Aldrich). In gel cleavage assays were performed in biological triplicates using 6 μM β-casein (Sigma-Aldrich) as a substrate. Samples were taken out at timepoints 0 min, 10 min, 20 min, 40 min, 60 min and 90 min, respectively, and immediately mixed with SDS-PAGE loading buffer and heated to 95 °C for 2 min. Samples were then analyzed by SDS-PAGE using 4–20% MiniProtean TGX gels (Bio-Rad) and the protein bands were visualized using SimplyBlue SafeStain (Thermo Scientific). Quantification of band intensities was performed subsequently using ImageJ (*46*).

### Fluorescence-based cleavage assay

Fluorescent cleavage assays were conducted using 0.8 μM hexFtsH or hexFtsH_E415Q_ (hexameric concentration) at 37°C in Buffer E supplemented with 2 mM ATP and casein fluorescein isothiocyanate (FITC-casein, Sigma-Aldrich). Cleavage of FITC-casein results in increased fluorescence thus allowing detection of proteolytic activity. The assay was run with varying concentrations of FITC-casein ranging from 1.4 μM to 60 μM. All components were incubated at 37 °C for 1 h before hexFtsH was added to start the cleavage reaction. Reactions were performed in biological triplicates, fluorescence was measured in 96-well plates (Corning) on a FLUOstar Optima plate reader (λ_ex_: 485 nm, λ_em_: 520 nm). The *K*_m_ was calculated by extracting the initial reaction rates and fitting the data to either the Michaelis-Menten equation (8) or an allosteric model (9) using GraphPad Prism 10.0.

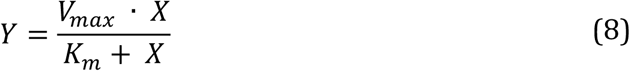

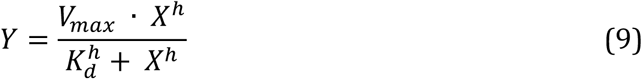

*v*_max_ and *k*_cat_ were calculated based on a standard curve using bovine trypsin cleavage of FITC-casein (**fig. S1A**).

### NADH coupled ATPase assay

Measurement of ATP activity was conducted using NADH-coupled ATP assays and β-casein (Sigma-Aldrich) as substrate. All reactions were carried out at 37 °C in buffer E together with 0.4 mM NADH (Merck), 20 U/mL lactate dehydrogenase (Roche) and an ATP regeneration system consisting of ATP in concentrations ranging between 8 – 1000 μM, 3 mM phosphoenolpyruvate (Roche) and 20 U/mL pyruvate kinase (Sigma-Aldrich). HexFtsH was added at a concentration of 0.8 μM (hexamer) as well as 4.8 μM β-casein, hexFtsH_E415Q_ was added at 0.1 μM and 1.0 μM β-casein. Reactions were carried out in biological triplicates and the consumption of NADH was measured at 340 nm in a 96-well plate (Greiner) using FLUOstar Optima plate reader. The ATPase rate was calculated using an NADH standard curve (**fig. S1B**) and fitted to a Michaelis Menten equation using GraphPad Prism 10.0.

### NMR-based ATP regeneration assay

ATPase activity was in addition assessed by 1D ^1^H NMR spectroscopy employing an ATP regeneration system directly in the NMR tube. The reaction was performed at 37 °C in D_2_O based buffer E with 5 mM ATP and the regeneration system consisting of 6 mM phosphoenolpyruvate, 20 U/mL pyruvate kinase. 100 μM β-casein was used to stimulate ATP consumption and after addition of 100 μM hexFtsH (monomer concentration), the reaction was incubated for 10 minutes at 37 °C before starting the experiments. 1D ^1^H NMR experiments were recorded for 5 minutes, and the data were processed in Topspin 4.4.0 (Bruker BioSpin) with line broadening set to 2. The increasing intensity of the pyruvate resonance was integrated using the same software. Resulting integrals were transformed into concentrations by a phosphoenolpyruvate standard curve (**fig. S1C**), as the increase of pyruvate is proportional to the decrease of phosphoenolpyruvate. The pyruvate increase was fitted to a simple linear regression model using GraphPad Prism 10.0 to extract kinetic rates.

### ATP single turnover assay

To characterize ATP activity under single-turnover conditions, hexFtsH was incubated with 10 mM ATP on ice for 10 minutes to minimize ATP hydrolysis. Unbound ATP was removed from the hexFtsH-ATP complex by gravity flow gel filtration (Nick columns, Cytiva), equilibrated with buffer D, in a cold room (∼8°C) adapting a previously reported protocol (*47*). After elution, the complex was snap frozen in liquid nitrogen. For the single turnover assay, samples were thawed on ice and mixed with prewarmed buffer E. The reaction was carried out at 37 °C and timepoints were taken out over time and immediately mixed with 300 mM HCl (1:1 volume) to stop the enzymatic reaction. Samples were kept on ice and centrifuged for 5 minutes, at 4 °C, 16.000 x g to remove possible precipitates. The supernatant was frozen in liquid nitrogen and kept at -80 °C until analysis upon which they were injected into a 6 mL Resource Q column (Cytiva) connected to an ÄKTA system (Cytiva) that was pre-equilibrated with 25 mM HEPES, pH 8. Chromatogram peaks of ATP at 260 nm were integrated using GraphPad Prism 10.0 and the data was fitted to a mono-exponential curve. The assay was run in biological triplicates.

### Mant-nucleotide titration assay

Nucleotide binding to hexFtsH was examined using the fluorescent nucleotide analogs 2’/3’-O-(N-Methyl-anthraniloyl)-ADP (mant-ADP) and 2’/3’-O-(N-Methyl-anthraniloyl)-ATPγS (mant-ATPγS) obtained from Jena Bioscience. Binding of the mant-nucleotides to proteins leads to an increase of the fluorescence of the mant group, allowing monitoring of the nucleotide binding event. To determine the affinity of Mant-nucleotides to hexFtsH, increasing concentrations of hexFtsH was titrated, 0.1–10 μM monomer concentration in 13 steps, to a fixed concentration (1 μM) of mant-ADP or mant-ATPγS, respectively, in buffer E. Fluorescence (λ_ex_: 355 nm, λ_em_: 435 nm) was measured using an Infinite 200 PRO Tecan plate reader in 96-well plates (Corning) at 37 °C. The reactions were run in biological triplicates, and the dissociation constant (*K*_d_) was determined by fitting the quadratic equation (10) of the normalized fluorescence *versus* hexFtsH concentration using GraphPad Prism 10.0.

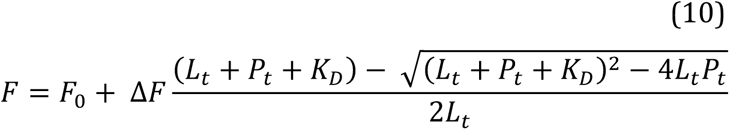

where *F*_0_ is the fluorescence of free m-nucleotide, *11F* = *F*_bound_ – *F*_0_, *L*_t_ is the total concentration of the mant-nucleotide, *P*_t_ is the total hexFtsH concentration, and *K*_d_ is the dissociation constant.

### Mant-nucleotide release assay

Nucleotide release from hexFtsH was performed using mant-nucleotides, 1.25 μM hexFtsH (monomer concentration) was incubated with 1.25 μM mant-ADP or mant-ATPγS in buffer E for 15 minutes at 37°C to ensure complete binding. To start the reaction, 80 μl of hexFtsH:mant-nucleotide complex was rapidly mixed with 20 μl of either 10 mM unlabeled ADP (Sigma-Aldrich), ATPγS (Jena Biocience), phosphate or a mix of both ADP and phosphate, in buffer E, using an automated injection system (Te-Inject) connected to an Infinite 200 PRO Tecan plate reader. Fluorescence was measured (λ_ex_: 355 nm, λ_em_: 435 nm) over time at 37°C in a 96-well plate (Corning). All reactions were carried out in biological triplicates. The final concentrations after mixing were 1 μM hexFtsH, 1 μM mant-nucleotide and 2 mM of the respective unlabeled nucleotides. The measured fluorescence was smoothed using a local average over five consecutive data points and subsequently normalized, *k*_off_ was determined by fitting the data to a mono-exponential function using GraphPad Prism 10.0.

### SEC-MALS

SEC-MALS experiments were performed using a Superdex Increase 200 10/300 GL column (GE Healthcare) on an Agilent 1260 HPLC Infinity II at room temperature. aFtsH, pFtsH and cFtsH were ran in buffer C. 2 mg/ml BSA (Sigma-Aldrich) was used as a calibration and to normalize the obtained data. Protein elution was monitored by an Agilent multi-wavelength absorbance detector, measuring at 280 nm and 254 nm, a Wyatt miniDAWN TREOS multiangle light scattering (MALS) detector and a Wyatt Optilab rEX differential refractive index (dRI) detector. The data were processed using the ASTRA 7.1.3 software (Wyatt Technology).

## Results & Discussion

### Isolated domains of cytoplasmic FtsH in solution

*E. coli* FtsH spans the inner membrane containing a small periplasmic domain and a larger cytoplasmic portion. The cytoplasmic domain represents the core part of the catalytic activity of FtsH consisting of the two functional domains, the AAA+-ATPase and the protease domain (**Fig. 1A**). To date, structural information about the functional domains of *E. coli* FtsH remains scares, as it is mainly based on a crystal structure of the AAA+ sub-domain (*11*), and low resolution cryo-EM structures of full-length FtsH in a larger complex together with regulators HflKC (*14*, *48*). To study the functional and structural details of FtsH using advanced solution NMR-spectroscopy, we therefore adopted a divide-and-conquer approach, facilitated by the domain organization of the FtsH cytoplasmic portion to create smaller subconstructs. Constructs of the isolated cytoplasmic domain (cFtsH), AAA+ domain (aFtsH) and protease domain (pFtsH) were prepared (**Fig. 1B**). We initially produced uniform labeled [*U*, ^2^H,^15^N] cFtsH and recorded a 2D ^15^N NMR spectrum. The resulting spectrum (**Fig. 1C**), is of decent quality but showed a large degree of varying peak intensities, suggesting different dynamic behavior, as well as partial signal overlap, which taken together prevents further sequence specific resonance assignment. On the other hand, the resulting ^15^N NMR spectra (**Fig. 1D** and **E**) of the isolated AAA+ domain and the isolated protease domain, indicate well-folded subdomains and are of notably higher quality compared to the larger construct. Although the peak intensities still vary to some extent for both the AAA+ domain, and particularly for the protease domain, pointing to potential oligomerization. Analysis of oligomeric states by SEC-MALS for the constructs (**fig. S2** and **table S4**) confirmed that all constructs mainly behave as monomers, with a ∼30-40 percentage of the AAA+ domain forming a dimeric species at room temperature under the used conditions. Based on a suite of 3D NMR through-bond experiments, we were able to assign the backbone resonances in a sequence-specific manner, resulting in ∼95% complete assignment for the AAA+ domain and ∼50% of the protease domain. Based on this resonance assignment, the secondary structure propensities in solution of the isolated domains were extracted from the secondary ^13^C chemical shifts (**Fig. 1F**) and compared to the secondary elements reported from the crystal structure of the AAA+ domain and the predicted AlphaFold3 structural model of the protease domain. The secondary elements show overall good agreement, confirming that the isolated domains adopt the same secondary structure in solution. The missing assignments for the AAA+ domain can mainly be attributed to residues in flexible loops and linker regions, which under the conditions used, pH 7.4 and elevated temperature of 310K, can be explained by enhanced amide-proton exchange rates in these unstructured regions (*49*). In contrast, we observed for the protease domain a large degree of unstructured elements in line with the structural model. This is also evidenced by a sub-set of high intensity signals observed in the ^15^N-NMR spectrum (**Fig. 1E**), impairing a full sequence specific resonance assignment. Therefore, we focused further our characterization mainly on the AAA+ domain.

**Figure 1.**
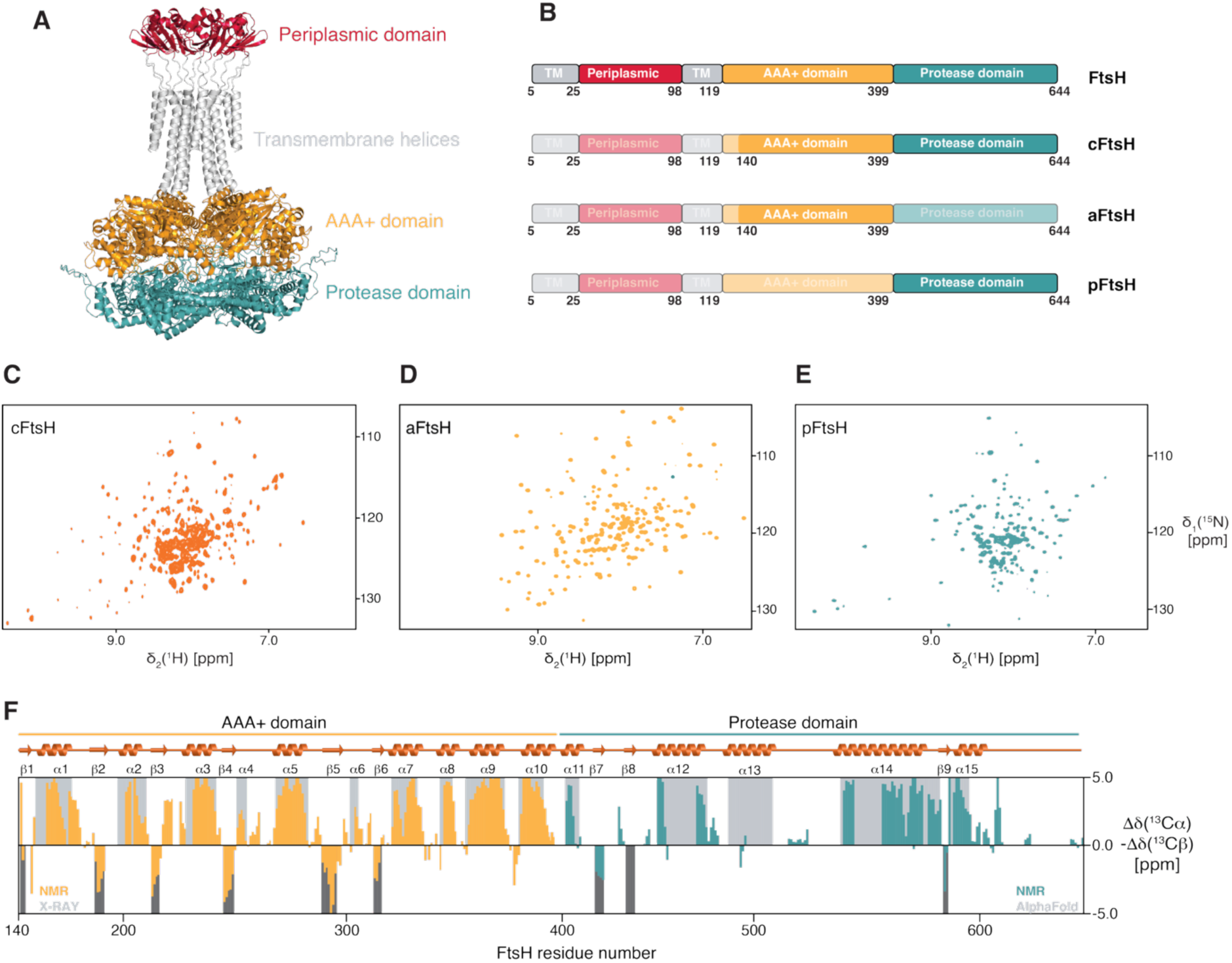
FtsH domain structure. **A)** Structural model of full-length *E. coli* FtsH obtained from Alphafold3 (*50*). **B)** Schematic representation of the FtsH functional domains and the constructs used in the present study. **C–E)** 2D ^15^N-NMRspectra of [*U*-^2^H,^15^N]-cFtsH (**C**), [*U*-^2^H,^15^N]-aFtsH (**D**) and [*U*-^2^H,^15^N]-pFtsH (**E**), respectively. **F)** Secondary structure elements of aFtsH and pFtsH derived from the secondary ^13^C chemical shifts (orange, blue) alongside the schematic representation of the secondary structure elements (grey shading) of the AAA+ domain derived from the crystal structure (PDB-ID: 1LV7) and of the protease domain from the AlphaFold3 prediction (AlphaFoldDB-ID: P0AAI3). Positive values indicate the presence of α-helical regions whereas negative values β-strand regions.

### Backbone dynamics of aFtsH reveal the interplay of its two sub-domains

To investigate the dynamical behavior of aFtsH, we recorded NMR relaxation experiments probing backbone dynamics across different timescales (*34*, *51*). Measuring ^15^N{^1^H}-NOE (hetNOE) and ^15^N longitudinal *R*_1_ relaxation rates, to obtain information on the fast motions on the pico-to nanosecond timescale, whereas the ^15^N transverse *R*_2_ relaxation rates, extracted from the *R*_1ρ_ rates, report on the micro- to millisecond timescale. Overall, we observed a stable hetNOE ratio profile on average 0.81 ± 0.1 and 0.79 ± 0.1 for 21.1 T and 18.8 T, respectively (**fig. S3**). Overall, the obtained values are well within the theoretical maximum value expected at 18.8 T (800 MHz ^1^H frequency) of 0.86 pointing to a stably folded protein whilst also ruling out experimental artifacts due to incomplete relaxation (*52*). Plotting the obtained values onto the AAA+-domain crystal structure (PDB-ID: 1LV7 (*11*)) reveals a stable fold for both sub-domains, whereas enhanced local flexibility can be observed around the two Walker motifs (e.g. G195–T202 and I247–D254) as well as the residues forming the substrate-threading pore around F228, suggesting inherent flexibility in the apo-state (**Fig. 2A**). Besides these mentioned regions, we also detected lower hetNOE values for the carboxy-terminal part of helix α1. This trend indicates the presence of local fast dynamics, which was also evident to some extent from the obtained secondary ^13^C chemical shifts, suggesting not a stable long helix but rather up to three helical segments (**Fig. 1F**). For the *R*_1_ relaxation rates, we observed a similar trend with enhanced rates for the regions mentioned above, experiencing fast dynamics albeit having average rates of 0.915 ± 0.12 s^-1^ and 0.933 ± 0.1 s^-1^, respectively (**fig. S3**). Using the program Tensor (*36*), we determined the rotation correlation time (τ_c_) yielding a value of ∼7.88 ns. The obtained value seemed on a first glance low considering that the aFtsH construct with its size of 26 kDa was expected to have a τ_c_ ∼15 ns (*53*). Nevertheless, considering that the ATPase domain contains two sub-domains, in line with earlier structural studies on FtsH from *Thermotoga maritima* (*54*, *55*), suggesting a large degree of inherent flexibility between the amino-terminal and carboxy-terminal part of the ATPase domain. This would match our observation of a possible partial decoupling of these two sub-domains leading to a lowered τ_c_.

**Figure 2.**
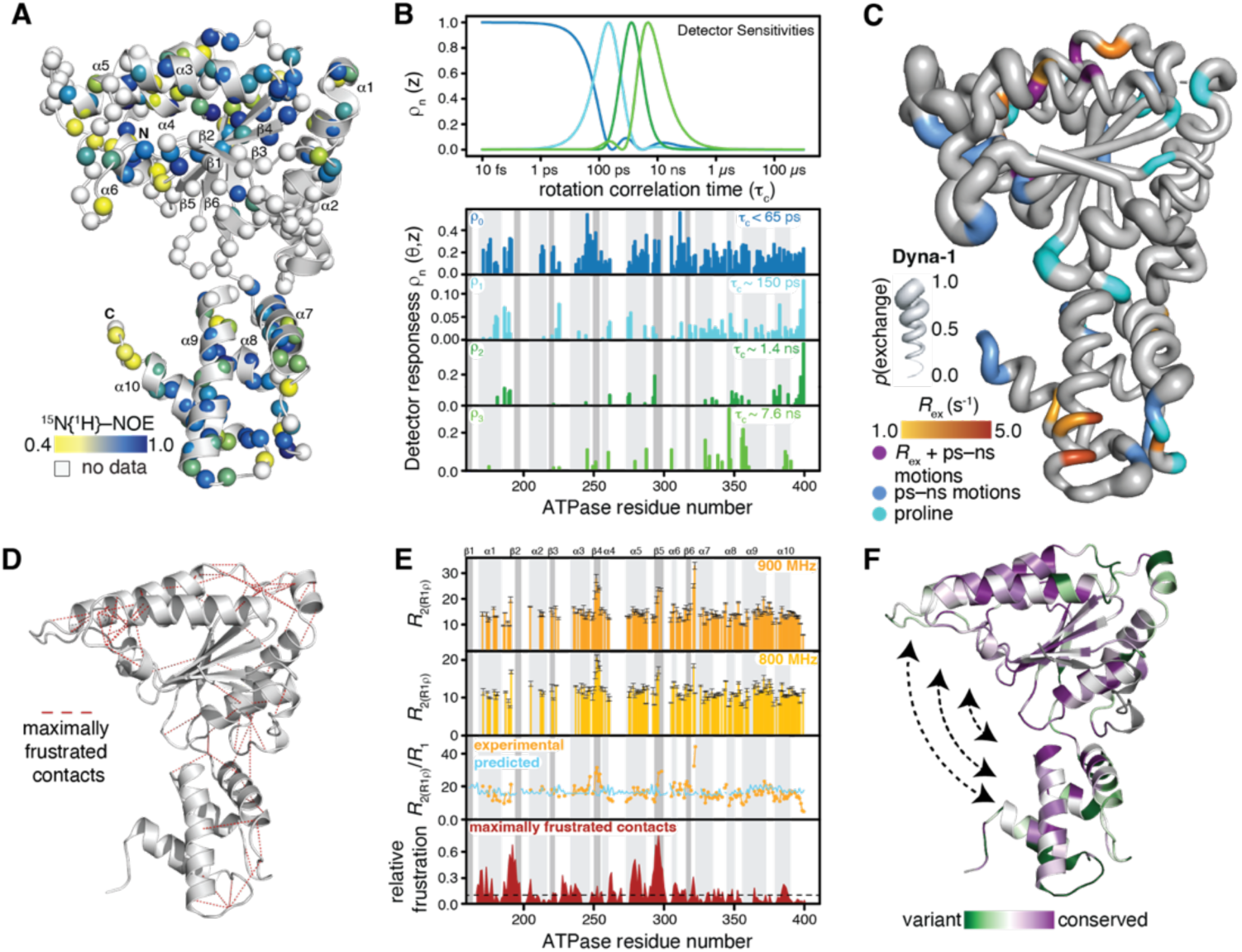
Dynamics of the FtsH-AAA+ domains indicate an allosteric coupling. **A)** Dynamics on the pico- to nanosecond timescale plotted on the AAA+ domain structure (PDB–ID: 1LV7). The amide moieties are depicted as spheres and the hetNOE values are indicated by the green to blue gradient. **B)** Analysis of the distribution of the inherent dynamics extracted from the NMR relaxation data using the detectors approach (*40*, *41*) (github.com/alsinnmr/pyDR). Sensitivity of the four distinct detectors, *ρ*_n_, to different time scales of motions (top). Detector responses to the four obtained detectors (bottom), reflecting amplitudes of motions in the time windows depicted on top. **C**) Dyna-1 (*60*) derived theoretical degree of micro- to millisecond exchange within the FtsH AAA+-domain indicated by the tube thickness. Experimentally derived chemical exchange contributions, *R*_ex_ as extracted by the Lipari-Szabo model free approach (*57*, *58*), indicative of micro- to millisecond motions are shown by the yellow to red gradient. Residues experiencing extensive motions on the pico- to nanosecond timescale, as derived from the extent of the hetNOE values (panel **A**), are indicated in blue for values below 0.65 and residues experiencing extensive motions on both timescales are highlighted in purple, whereas prolines are shown in cyan. **D**) Structural frustration analysis was calculated for the AAA+ domain structure (PDB-ID: 1LV7) *via* the Frustratometer webserver (*61*). Highly frustrated interactions are indicated by the red lines. **E**) Analysis of the micro- to millisecond motions. Top panels show the *R*_2(R1ρ)_-rates derived from the measured *R*_1ρ_ relaxation rates at 21.1 T (orange) and 18.8 T (yellow). The experimentally obtained *R*_2(R1ρ)_/*R*_1_ ratio was compared to the theoretical values assuming rigid tumbling obtained *via* HYDRONMR (*39*). The density of the maximally frustrated contacts in the AAA+ domain was obtained *via* the Frustratometer webserver (*61*). **F**) Sequence conservation analysis using ConSurf (*62*, *63*). The used gradient from green (variable) to purple (conserved) is indicated. Arrows indicate the observed flexible arrangements between the two sub-domains.

In a next step we used the classical Lipari-Szabo model free approach to quantitate the motions within aFtsH (*56–58*). In line with the initial analysis of the measured relaxation parameters, we obtained a generally flat profile with an average value of 0.83 ± 0.06 for the generalized order parameter S^2^ (**fig. S4**). The obtained value is indicative of a stable protein fold in respect to the fast motions of the N–H vector. To further quantify the inherent motions, we next used the Detectors approach introduced by Smith and co-workers (*40*, *41*). In this approach, the relaxation analysis is not fitted to a single process at one timescale, which is most commonly done in relaxation data analysis (*59*), but rather reports on the amplitudes of motions in different time windows (referred to as detectors). Thus, the spin relaxation is modeled using a combined dynamical process, taking correlation times over the whole nano– to millisecond time window into account. This is done under the assumption of a distribution of correlation times *θ* (z), on a logarithmic scale, with z = log_10_(τ_c_/s). Using four detectors in the analysis, reporting on the distribution of motions over the range of the correlation time, we obtained the best fit to the experimental relaxation data (**Fig. 2B** and **fig. S5**). The obtained response to these detectors (indicating their respective amplitudes), suggests the largest part is related to enhanced picosecond motions as evidenced by the two detector windows < 65 ps and 150 ps (**Fig. 2B**). In the nanosecond timescale, the extent of the motions is greatly diminished, showing only distinct parts such as the carboxy terminus in the 1.4 ns window and the region around helix α8 for the 7.6 ns window (**Fig. 2B**). Besides the terminus, we also observe enhanced motions in the vicinity to the substrate threading pore as well as within residues involved in ATP hydrolysis.

As we observed extensive line-broadening for several residues at the hinge region between the two sub-domains as well as of parts the pore threading region, thus limiting our relaxation analysis for these regions severely, we employed the Dyna-1 approach (*60*) to obtain a more complete picture. Dyna-1 uses deep-learning models to predict missing sequence-specific assignments but has also shown the remarkable power to predict conformational exchange as measured by NMR relaxation (*60*). By this approach, we correlated our measured relaxation parameters with the predicted ones (**Fig. 2C**). Overall, we observe excellent agreement, as the regions not assignable in particular show a large degree of predicted conformational exchange on the micro- to millisecond timescale, as well as enhanced pico- to nanosecond motions.

Next, we focused our attention on the micro- and millisecond dynamics and their potential origin (**Fig. 2D, E**). We first characterized the obtained *R*_2(R1ρ)_ relaxation rates, yielding averages of 14.6 ± 3.4 s^-1^ and 11.8 ± 2.3 s^-1^ for 21.1 T and 18.8 T respectively (**fig. S3**). Mainly three regions showed enhanced *R*_2(R1ρ)_ rates, namely the Walker B motif I247–D254, the β_5_-strand as well as the linker between the two subdomains connecting strand β_6_ and helix α_7_ (**Fig. 2E**). To substantiate these observations further, we next compared the experimentally derived*R*_2(R1ρ)_/*R*_1_ ratios to the theoretical ones obtained from HYDRONMR (*39*), that assume anisotropic rigid tumbling for the molecule. Therefore, we presumed that the direct comparison would enable us to extract the experimentally determined conformational exchange contributions in more detail. Whereas we observed a global flat profile for the theoretical values, our experimental data indicate enhanced dynamics in the Walker B motif alongside the interdomain linker as already indicated by the enhanced *R*_2(R1ρ)_ rates discussed above.

Finally, we aimed to identify the origin of these conformational exchange contributions by characterizing the local frustration of the AAA+ domain (*61*). Remarkably, we observed in several already discussed regions, highly structurally frustrated contacts (**Fig. 2 D, E**): the pore threading loop, in the vicinity of the ATP binding site as well as the linker between the two sub-domains. Interestingly, we also detected a cluster of frustrated contacts at a possible hinge-region that might be involved in the substrate threading motion within the catalytically active hexamer. We were intrigued by this connection and wondered how much of these potentially functionally important dynamical properties are encoded already at the amino-acid level, and thus decided to analyze the protein conservation using ConSurf (*62*, *63*). Remarkably, we observed that the linker region between the two domains is highly conserved (**Fig. 2F**). Given the fact that these residues also comprise parts of the ATP-binding site, this illustrates that the observed flexibility in this area is likely directly modulated by the different nucleotide-bound states. In addition, large parts of the structural elements are highly conserved, whereas in contrast the pore threading loop shows a larger degree of variance on the sequence, which might possibly reflect partially differing substrate pools in different organism (*64*). Even though an aromatic residue is common as the key component in the substrate-threading loop, for the FtsH-related AAA+ proteins it is directly adjacent to another hydrophobic residue (e.g. valine or methionine). This residue is presumed to increase the hydrophobicity around the encountered substrate and thus directly reflects their respective pool of substrates (*65*, *66*).

### Methyl group assignment and side-chain dynamics of cytoplasmic FtsH

Based on the observations of an intricate coupling within the sub-domains of the ATPase part of the FtsH cytoplasmic domain, we wondered if we could extend this analysis further towards the whole cytoplasmic domain. Initial tests with this construct indicated that the quality of the ^15^N-NMR spectrum (**Fig. 1C**) is not sufficient. Therefore, we turned our attention to ^13^C-methyl NMR spectra, which show increased sensitivity, making them particularly useful for large proteins and their complexes (*67–69*). Based on the almost complete backbone and side-chain assignment of the AAA+ domain together with the partial assignment of the protease domain, we could already transfer a large extent of sequence-specific resonance assignments to separately expressed Met/Ile-cFtsH, Ala-cFtsH, and Val,Leu*^proS-^*cFtsH samples (**Fig. 3A**). These initial assignments were further substantiated and confirmed by measuring methyl-methyl NOESY experiments, leading to an extent of the methyl group assignment in total of 100% for the ATPase domain. On the basis of this assignment, we observed for a subset of resonances uncharacteristic features. We observed an unusual ^1^H chemical shift for M285, indicative for being bound to aromatics (*70*), suggesting either the formation of a sulfur-ν−interaction or a methyl-ν−interaction between the M285 and adjacent F316 (**Fig. 3B**), which has been shown to be an important stabilizer of protein interactions (*71*, *72*). Additionally, the analysis of its ^13^C-chemical shift further points to a functional relevance of this residue as this suggests it to be in the *trans*-state in solution in contrast to the crystal structure (**Fig. 3B**). As only about ∼9 % of methionine residues are found in the *trans*-state (*73*), together with the notion that the conversion of sidechain rotameric states is commonly observed in regions involved in allosteric rearrangements (*74*), this highlights the potential relevance of this residue which is likely also reflected by its high degree of sequence conservation (**Fig 2F**). Besides this interaction with an aromatic moiety, we also observed Hβ chemical shifts for two alanines in helix α_11_, namely A371, A376, as well as for A298 situated in strand β_4_, which suggest a close interaction with aromatic residues, e.g. F375, F388 and F316, respectively.

**Figure 3.**
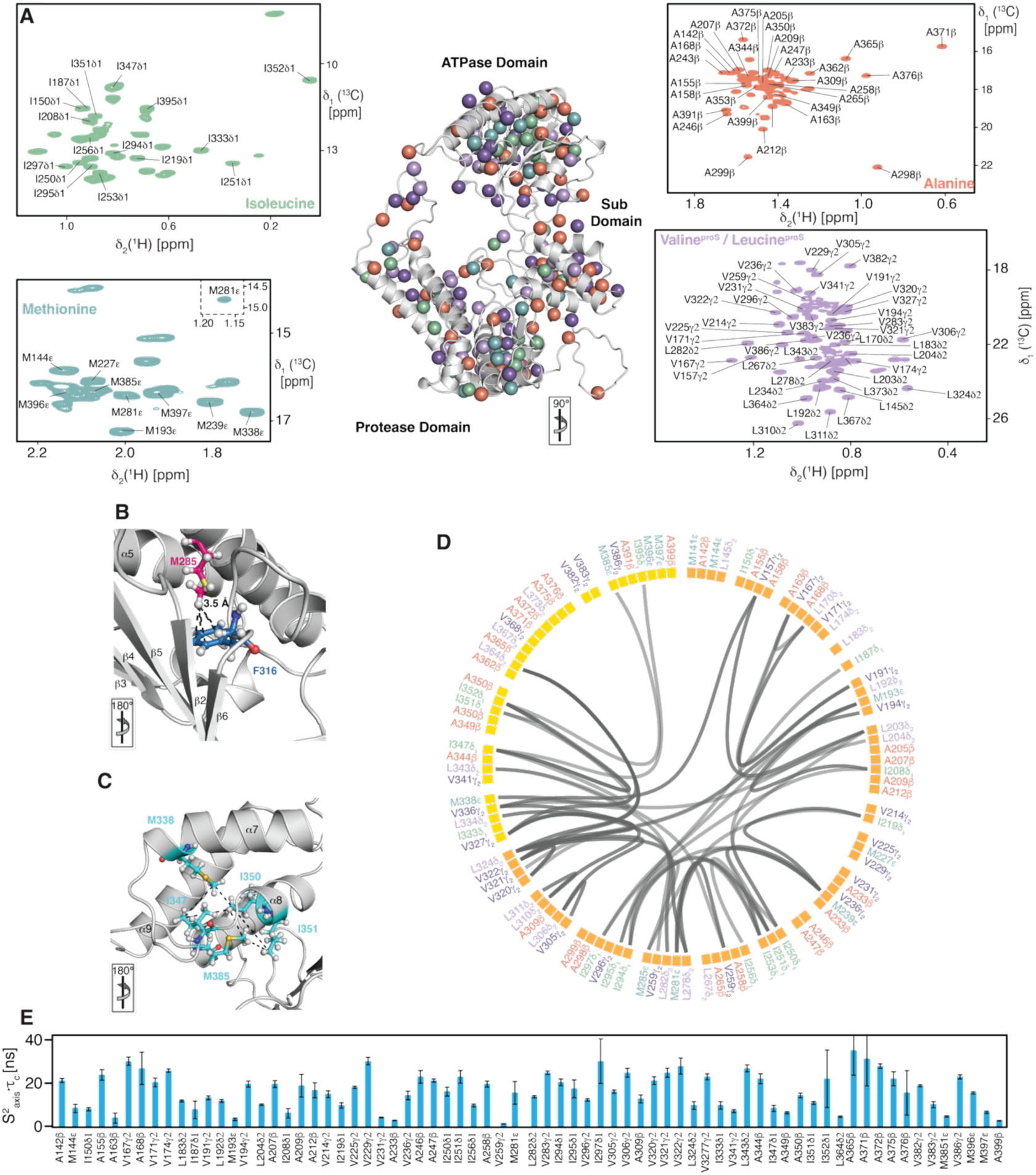
Methyl group interplay in the cytoplasmic domain and resulting side-chain dynamics. **A)** Distribution and sequence specific resonance assignment of the methyl groups of methionine (cyan), alanine (orange), isoleucine (green) as well as the *proS* methyl groups of valine (light purple) and leucine (purple). **B**) Close-up onto the methionine-phenylalanine motif stabilizing the ATPase sub-domain. **C**) Characteristic NOE network plotted on the AlphaFold3 prediction (AlphaFoldDB-ID: P0AAI3) of the FtsH cytoplasmic domain. In panels **A–C** is the rotation relative to Figure 2A indicated. **D**) Flareplot visualization of the methyl-methyl NOE network within the ATPase domain detected for the three individually labelled cFtsH samples illustrating the connectivity between the different sub-domains of the ATPase domain. **E**) Local methyl group dynamics on the pico- to nanosecond timescale probed by methyl single quantum (SQ) and triple quantum (TQ) relaxation experiments showing the product of the local order parameter and the overall tumbling constant, S^2^_axis_ • τ_c_.

Besides guiding the assignments, the NOESY experiments also facilitated the identification of crucial residues linking the different sub-domains within the cytoplasmic domain together (**Fig. 3D**). Using the three different samples already mentioned above, we were able to derive several methyl-methyl NOEs indicating general stabilization of the two sub-domains of the ATPase domain (**Fig. 3D**). This analysis also enabled us to reveal a network between methyl-bearing amino acids situated in the amino-terminal loop-region and α_2_, showing connections to helix α_7_ in the other sub-domain (**Fig. 3E**). Comparing these regions to the backbone relaxation analysis (**Fig. 2**), revealed enhanced inherent dynamics in these regions as discussed earlier. This further highlights the NOE network in modulating the dynamics between the two sub-domains and potentially being involved in allosteric signaling.

Having established the methyl NOE network within the larger construct, we next focused on characterizing the dynamics of the methyl groups. In a first step, we determined the product of the side-chain order parameter and the correlation time of the overall molecular tumbling (S^2^_axis_ • τ_c_), reporting on the extent of the amplitude of motions on the pico- to nanosecond timescale (*43*, *44*). The obtained values showed a maximum of ∼30–35 ns in line with the size of the entire cytoplasmic domain of 55 kDa. This potentially indicates, in contrast to the backbone relaxation properties reported for the isolated ATPase domain, a certain degree of rigidification of this larger construct. Overall, the distribution of the values indicates some inherent flexibility particularly in loop regions and the sub-domain interface, which for the isolated ATPase domain was already captured in the detailed backbone relaxation analysis discussed above, but also provides some initial insight into the dynamics of the protease domain (**Fig. 3F** and **fig. S6)**.

As the initial ^13^C-NMR spectra of the different methyl-labelled cFtsH samples showed some indication of specific line-broadening, which can potentially be attributed to conformational exchange processes, we next used a multiple quantum (MQ) Carr-Purcell-Meiboom-Gill (CPMG) relaxation dispersion experiment (*45*). We quantified the exchange-induced broadening effects (depicted as 11*R*_2,eff_) on the micro- to millisecond timescale by measuring the difference at two different fields (33 and 1000 Hz) and at two magnetic fields 16.4 T and 18.8 T (**fig. S6**). Albeit the vast majority of the residues showed an absence of broadening effects, and thus methyl dynamics on the microsecond timescale, we observed dispersion for a sub-set of resonances. This includes dispersion at the interface between the two sub-domains, as well as in the region linking the ATPase and the protease domain. Taken together a picture emerges, where in the cytoplasmic part of FtsH, all three sub-domains are connected in a highly dynamic yet controlled manner, highlighting the underlying allosteric signaling from the site of ATP consumption to the active site of the protease domain.

### Mapping ATP binding onto the ATPase domain

After having established the dynamics and the interplay between the individual subdomains comprising the cytoplasmic domain, we turned our attention to the interaction with ATP of the AAA+ domain. Structurally, FtsH contains two Walker motifs required for the ATP turnover, which are situated at the interface between two FtsH protomers within the active hexamer. Thus, even though ATP conversion requires the full hexamer, ATP binding can already be assessed by using the isolated ATPase domain. To this end we used a ^13^C-labelled aFtsH to monitor the resulting chemical shift and/or intensity changes on the methyl groups (**Fig. 4A–D**). With increasing ATP concentration, we observed both chemical shift perturbations as well as signal attenuations for a subset of residues (**Fig. 4A).** These observed effects indicate that on a chemical shift timescale, an interaction on the fast to intermediate regime was observed pointing to a dissociation constant in the low micromolar range (*75*). Mapping subsequently the observed effect onto the AAA+-domain structure (PDB-ID: 1LV7) showed good agreement with the positioning of the ATP in the crystal structures of related bacterial organisms (*54*, *76*) (**Fig. 4B** and **D**). Next, we wondered if we could elucidate the underlying allosteric signaling also towards the protease domain and thus repeated the ATP titration using a MI-cFtsH sample (**Fig. 4E, F).** To our surprise we observed a less pronounced effect in comparison to the isolated AAA+ domain, suggesting in agreement with our methyl group dynamics results a rigidifying effect of the protease domain which potentially dampens the extent of inherent dynamics of the AAA+-domain. Further, the absence of direct signaling from the AAA+-domain to the protease domain within one polypeptide chain suggest the need of cooperativity between the individual subunits of the FtsH hexamer. This interpretation is in full agreement with the need of a hexameric FtsH to ensure a functional ATPase cycle with individual ATP molecules bound at the interfaces of the respective subunits.

**Figure 4.**
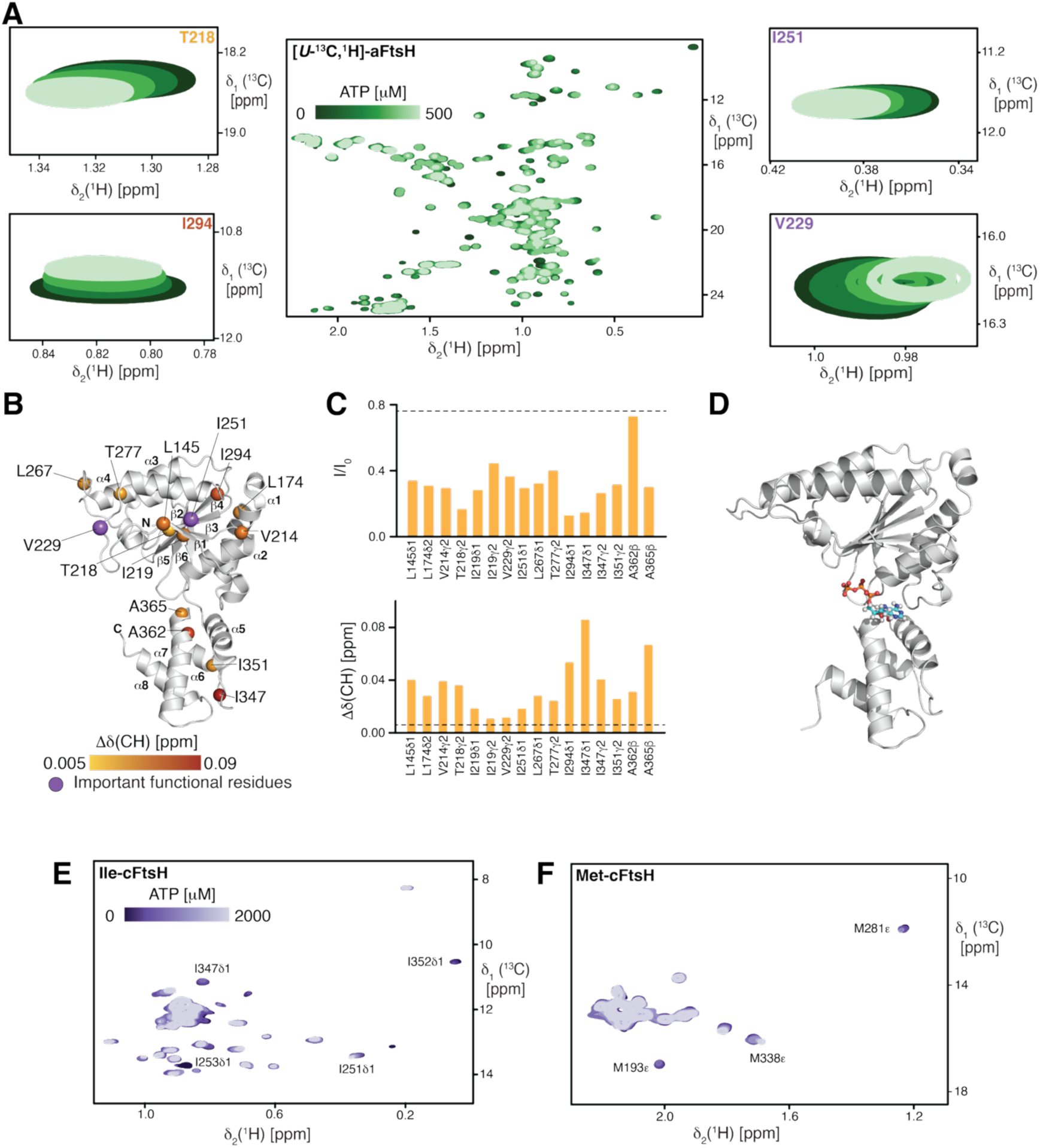
NMR titrations reveal ATP binding site and stabilizing effect on AAA+ domain. **A)** Chemical shift perturbations (CSPs) observed for the methyl groups of the ATPase domain upon ATP addition. **B)** Chemical shift perturbations plotted onto the AAA+ domain structure (PDB–ID: 1LV7). CSPs are indicated by the yellow to red gradient and catalytically important residues indicated in purple, V229 of the pore loop motif and I251 of the Walker B motif. **C)** Significant signal attenuations (top) and chemical shift perturbations (bottom) plotted against the residue number. Broken lines represent a significance level in both plots. D) ATP docked into the AAA+ domain structure (PDB–ID: 1LV7) based on aligning it to ATP-bound *A. aeolicus* structure (PDB–ID: 8VWC) **E, F)** Isoleucine region **(E)** and methionine region **(F)** of a ^13^C-NMR spectrum of MI-cFtsH in the absence and presence of increasing amounts of ATP as indicated.

### hexFtsH as a soluble proxy for catalytically active FtsH

To investigate the functional details of FtsH in more detail, we looked to design a soluble hexameric FtsH construct. To this end we followed a previously outlined approach (*15*, *77*), by replacing the amino-terminal part of FtsH with a hexamerization peptide (ccHex). At the sequence level, this peptide has multiple repeats of the largely hydrophobic ELKAIA sequence-motif (**Fig. 5A–C**). To confirm the hexameric state of this new construct termed hexFtsH, we analyzed size-exclusion chromatography elution profiles, which proved to be consistent with the expected molecular weight of the hexameric state (360 kDa; **Fig. 5D**). In a next step, we confirmed the proteolytic activity of hexFtsH by in-gel cleavage assay, using β-casein as substrate (**Fig. 5E–G**). The results confirm that this artificial hexamer is indeed active, and almost completely degrade all β-casein after 90 minutes in the presence of ATP. Conducting the cleavage assay by replacing ATP with a non- or very slowly hydrolysable ATP analogue, ATPγS, resulted in almost no cleavage of β-casein (**Fig. 5D–G**). We further characterized hexFtsH by creating a proteolytically inactive mutant (hexFtsH_E415Q_), where the mandatory glutamic acid within the zinc binding motif (H**E**AGH) has been changed into a glutamine (*78*). In agreement, hexFtsH_E415Q_ was unable to cleave β-casein, in the presence of either ATP or ATPγS (**Fig. 5D–G**). In order to impair ATP hydrolysis, we also constructed a mutant in the Walker A nucleotide binding motif, hexFtsH_K201N_, which has been shown previously to significantly reduce ATPase activity and abolish proteolytic activity (*78*). In our hands, hexFtsH_K201N_ was also unable to cleave β-casein in the presence of either ATP or ATPγS (**Fig. 5D–G**). Comparing the catalytic efficiency under the different tested conditions revealed that both the E415Q-mutant, or the presence of ATPγS reduced it by a factor of ∼5 to 21% compared to hexFtsH in the presence of ATP (**Fig. 5G**). The other conditions had an even more drastic effect leading to a reduction of 8–14%, rendering the protease almost completely inactive (**Fig. 5G**).

**Figure 5.**
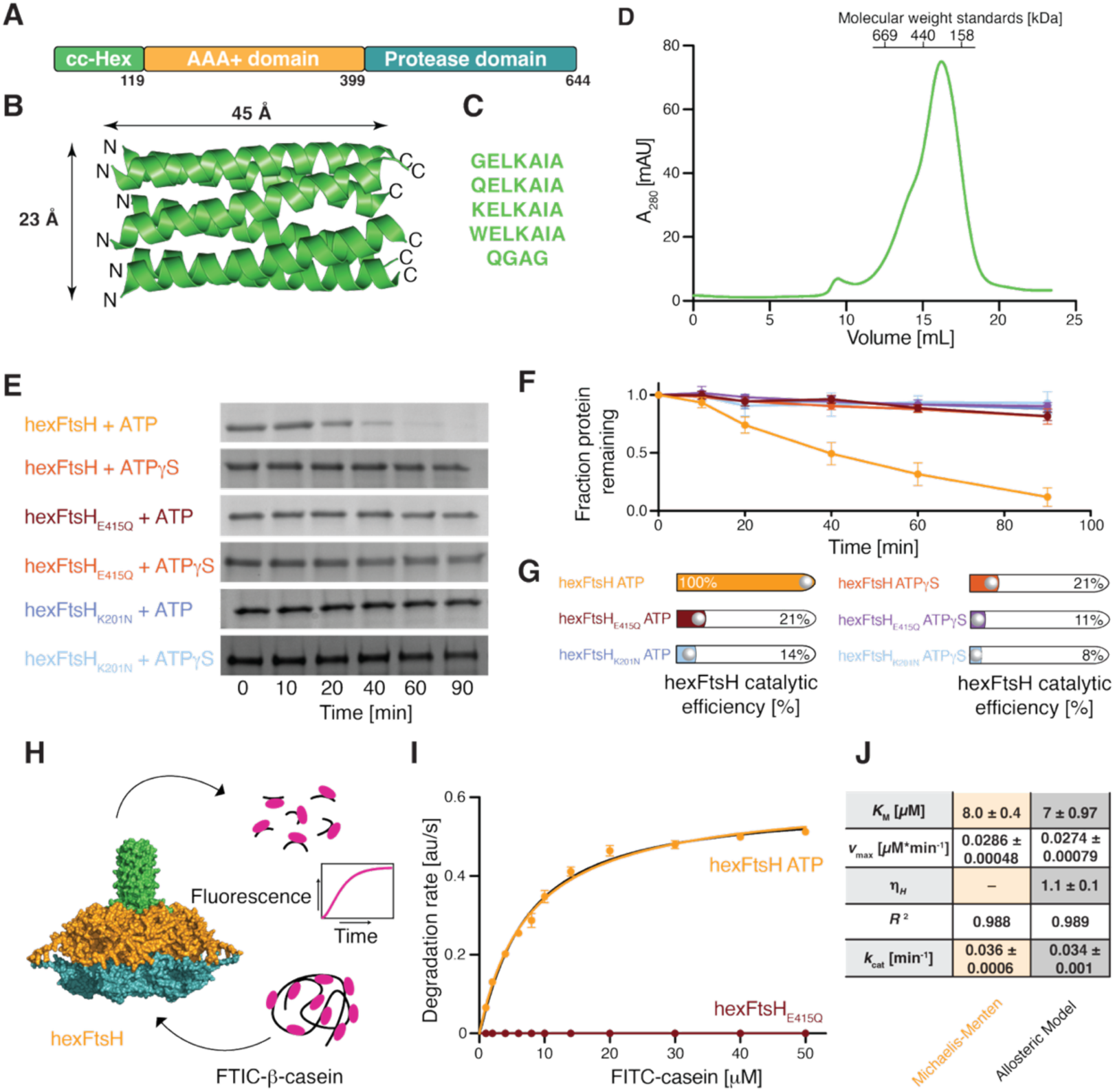
Using the hexFtsH-variant as a proxy for FtsH-function. **A)** Schematic representation of hexFtsH an amino-terminal fusion of FtsH to the hexamerization domain cc-Hex. **B**) Crystal structure of the cc-hexameric coiled-coil (PDB-ID: 3R3K) indicating the dimensions of the resulting hexamer. **C**) Sequence of the cc-Hex peptide, highlighting its inherent sequence repeats. **D**) Size-exclusion chromatogram obtained for hexFtsH. The molecular weights of a standard calibration curve are indicated at the top. **E**) Proteolytic cleavage of β-casein by different hexFtsH variants as indicated using either ATP or its slowly hydrolyzing analogue ATPγS. In-gel cleavage assays were run in biological triplicates yielding similar results. Uncropped gels and controls are shown in **figs**. **S7** and **S8**. **F**) Analysis of the in-gel cleavage assays depicted in panel **E**. **G**) Comparison of the proteolytic efficiency of hexFtsH towards β-casein for the different set-ups tested. Cleavage is expressed as percentage of β-casein cleaved after 90 minutes of incubation with the respective hexFtsH variants. Bar colors correspond to the colors assigned to each variant in panel **E**. **H**) Schematics of the employed fluorescence cleavage assay using the self-quenching fluorescein-5-isothiocyanate-(FITC)-casein as a substrate. **I**) Fluorescence-based assay reporting on the cleavage activity of hexFtsH (orange) and hexFtsH_E415Q_ (dark-red) employing variable concentrations of the FITC-casein substrate. Solid lines represent least-square fits to either the Michaelis-Menten degradation kinetics (orange) or to an allosteric model (black). Data is representative of three replicative experiments. **J**) Extracted kinetic parameters for both models obtained from the data shown in panel **I**.

To obtain more insight into the kinetics of the proteolytic reaction we next used a fluorescence-based approach employing self-quenching substrate-analog fluorescein-5-isothiocyanate-casein (FITC-casein; **Fig. 5H**). Using varying concentrations of the FITC-casein substrate, we extracted kinetic parameters for hexFtsH and hexFtsH_E415Q_ (**Fig. 5 I, J**). Whereas we did not observe any cleavage for the catalytically inactive E415Q-mutant, we could directly extract the *K*_m_ value of 8 ± 0.4 µM for hexFtsH using the Michaelis-Menten approach. We also tested if the obtained data would be better fitted to an allosteric model, given the hexameric state of FtsH (**Fig. 5 I, J**). The obtained *K*_m_-value of 7.0 ± 0.97 µM is almost identical to the Michaelis-Menten model, and even more importantly the Hill-coefficient of 1.1 ± 0.1 suggests the absence of allosteric coupling for the proteolysis reaction. Overall, the obtained kinetic parameters are in good agreement with previous data reported for the wild-type FtsH (*79–81*), highlighting the relevance of using the hexFtsH construct to assess FtsH function in detail.

### Assessing the functional ATP cycle

To investigate the ATP cycle for hexFtsH, we set out to characterize the length of the catalytic cycle by the recently introduced *in-cyclo* NMR approach (*82*). By using an ATP regeneration system, the consumed ATP is replenished which results in steady-state conditions (**Fig. 6A**). To this end, the enzyme pyruvate kinase (PK) is added which converts phosphoenolpyruvate (PEP) and ADP back to ATP, ensuring constant ATP levels throughout the experiment (**Fig. 6D**). By monitoring the ATP, PEP, and resulting pyruvate concentrations by ^1^H-NMR shows a linear depletion of PEP alongside the production of pyruvate, both at the same rate (**Fig. 6D**). As these observed changes are directly coupled in a stoichiometric manner to the ATP hydrolysis rate, the ATP hydrolysis rate can be directly determined from the increase in pyruvate concentration (**Fig. 6B**). Using this approach, we observed, in the presence of 100 µM hexFtsH (monomer) and substrate, an ATP consumption of 0.12 ± 0.093 mM min^-1^, which corresponds to a hydrolysis rate of *k*_hydr_ of 1.16 ± 0.06 min^-1^ per active site (**Fig. 6C**). The inverse length of the obtained rate, *T* = 1/*k*_hydr_ = 51.7 ± 2.7 s is thus the average length of the catalytic cycle.

**Figure 6.**
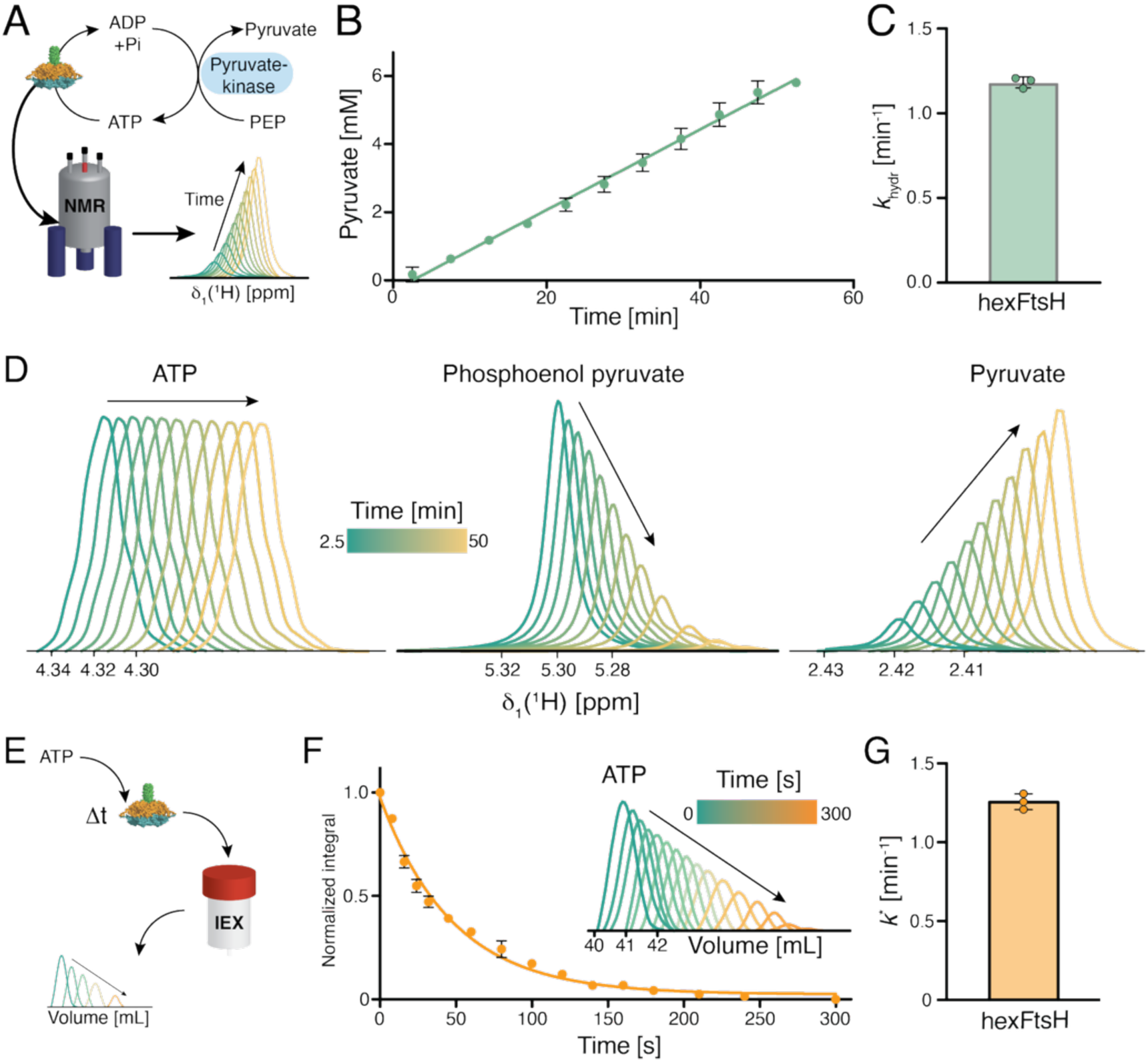
Assessing the functional cycle of ATP turnover by *in-cyclo* NMR and chromatography. **A)** Set-up of the *in-cyclo* NMR experiment adapted from Mas and Hiller (*82*). The ATP regeneration reaction using phosphoenolpyruvate (PEP) and the enzyme pyruvate kinase (PK) depicted, will run directly within the NMR spectrometer allowing access to the kinetic parameters directly by integrating ^1^H-NMR signals. **B)** Quantification of the pyruvate (green) concentration extracted from 1D ^1^H-NMR spectra shown in (**D**). Error bars indicate the standard deviation of three independent experiments. The solid line shows a linear fit to extract the ATP consumption rate. **C**) Molecular hydrolysis rate (*k*_hydr_) determined by the *in-cyclo* approach. Data points represent three independent experiments, and the error bars indicate the standard deviation. **D**) Series of 1D ^1^H NMR spectra of the ATP indole H8 resonance (left), the PEP resonance (middle) and the pyruvate resonance (right), measured in 5 min steps as indicated by the yellow to green gradient. **E**) Set-up of the adapted (*47*) ion exchange chromatography (IEX)-based assay to determine the hydrolysis rate of ATP, *k*\*. **F**) Normalized decay of ATP over the time course of the experiments as single point measurements at the indicated time points. Data points show the mean of three independent experiments whereas the error bars indicate the standard deviation. Inset shows the resulting chromatograms focusing on the ATP signal. **G**) Molecular hydrolysis rate (*k*_*_) determined by the chromatography approach. Data points represent three independent experiments, and the error bars indicate the standard deviation.

To assess the catalytic hydrolysis rate of ATP to ADP independently, we benchmarked the already discussed *k*_hydr_ rate with standard single-turnover experiments (*47*, *83*). In this set-up (**Fig. 6E**), the hexFtsH:ATP complex is initially formed at 4°C, assuming that the hydrolysis rate can be neglected under these conditions. Unbound nucleotides were removed by gel filtration, and the samples were flash frozen. The complex was then incubated at 37°C in the presence of a substrate and at distinct timepoints, the reaction was quenched by the addition of HCl. This leads to unfolding of the protein, thereby stopping the reaction immediately and releasing all educts and products into the solution. The samples were subsequently analyzed by anion exchange chromatography to determine the change in ATP concentration. The obtained data was fitted to a mono exponential decay resulting in a hydrolysis rate *k*\* of 1.3 ± 0.001 min^-1^ (**Fig. 6F, G**). This value is in excellent agreement with the rate determined by the *in-cyclo* NMR approach as the single turnover experiments commonly yield slightly larger hydrolysis rates compared to steady-state experiments (*82*).

### Assessing the nucleotide binding to FtsH

In order to assess the kinetic rates underlying nucleotide binding we adapted a standard fluorescence-based assay commonly used in the ATPase field (*84*, *85*), that on the one hand provides access to nucleotide binding, and on the other hand to nucleotide lifetimes by displacement (**Fig. 7A**). Using the fluorescently labelled ADP derivative, 2’-(3’-)-O-(N’-Methanylthraniloyl) adenosine-5’-diphosphate (mant-ADP) we initially measured the dissociation constants using increasing concentrations of hexFtsH and hexFtsH_E415Q_, respectively (**Fig. 7B**). We obtained a *K*_d_ of 1.4 ± 0.17 µM for hexFtsH compared to an almost identical *K*_d_ of 1.5 ± 0.07 for hexFtsH_E415Q_ (**Fig. 7C**). As we cannot obtain the values for ATP binding due to its fast turnover, we resorted to use its slowly hydrolysable analogue mant-ATPγS to assess nucleotide triphosphate binding. Whereas the experiment for hexFtsH resulted in a *K*_d_ of 1.0 ± 0.07 µM, we observed a slightly lower *K*_d_ for hexFtsH_E415Q_ with 0.57 ± 0.06 (**Fig. 7C**), in agreement with the estimated dissociation constants derived from the NMR titration experiments discussed above **(Fig. 4)**. Although the obtained values are almost identical, statistical analysis revealed that the values for nucleotide triphosphates were significantly lower compared to the diphosphates, potentially reflecting the underlying mechanism required for nucleotide turnover. To also get a more detailed picture over the lifetimes of the different bound nucleotides, we next performed a displacement experiment assessing nucleotide release by real-time fluorescence using an excess of non-fluorescent nucleotide (**Fig. 7A**). For hexFtsH, we observed almost identical *k*_off_ rates for ADP with 0.28 ± 0.03 min^-1^ and for ATPγS with 0.31 ± 0.05 min^-1^, respectively. In contrast, we observed a notable difference for hexFtsH_E415Q_ with 0.31 ± 0.04 min^-1^ for ADP compared to 0.62 ± 0.05 min^-1^ for ATPγS, suggesting that the presence of a non-functional catalytic site might accelerate the ATP turnover by limiting the ATP lifetime, a feature that is not possible in hexFtsH_E415Q_. The final step of the ATP catalytic cycle, ADP release can be influenced by both products, ADP and phosphate. Therefore, we also probed the displacement in the presence of saturating amounts of phosphate to discern which of the two products would be the rate limiting factor. Repeating the ADP displacement titration in the presence of phosphate resulted in an increased *k*_off_-rate of 0.48 ± 0.03. This value is notably higher than the rate determined for ADP alone by 0.28 ± 0.03 min^-1^ and would therefore indicate that the phosphate release is the rate limiting step within the FtsH ATP cycle. Taking into account the increased off-rate in the presence of phosphate, we wondered how the ratio of the different components is under physiological conditions. According to reported values in the literature the ratio of ATP:ADP between 1:0.8 and 1:0.3 with ATP concentrations ranging between 1–3 mM (*86*, *87*). Inorganic phosphate on the other hand is in the cytosol available at concentrations up to 10 mM, ensuring efficient replenishing of the nucleotide triphosphate pool in the bacterial cell (*88*, *89*). Thus, a picture emerges, where the presence of the phosphate speeds up the product release of the ATPase cycle under native like conditions, and relieving inherent limitations induced by the presence of a functional protease domain.

**Figure 7.**
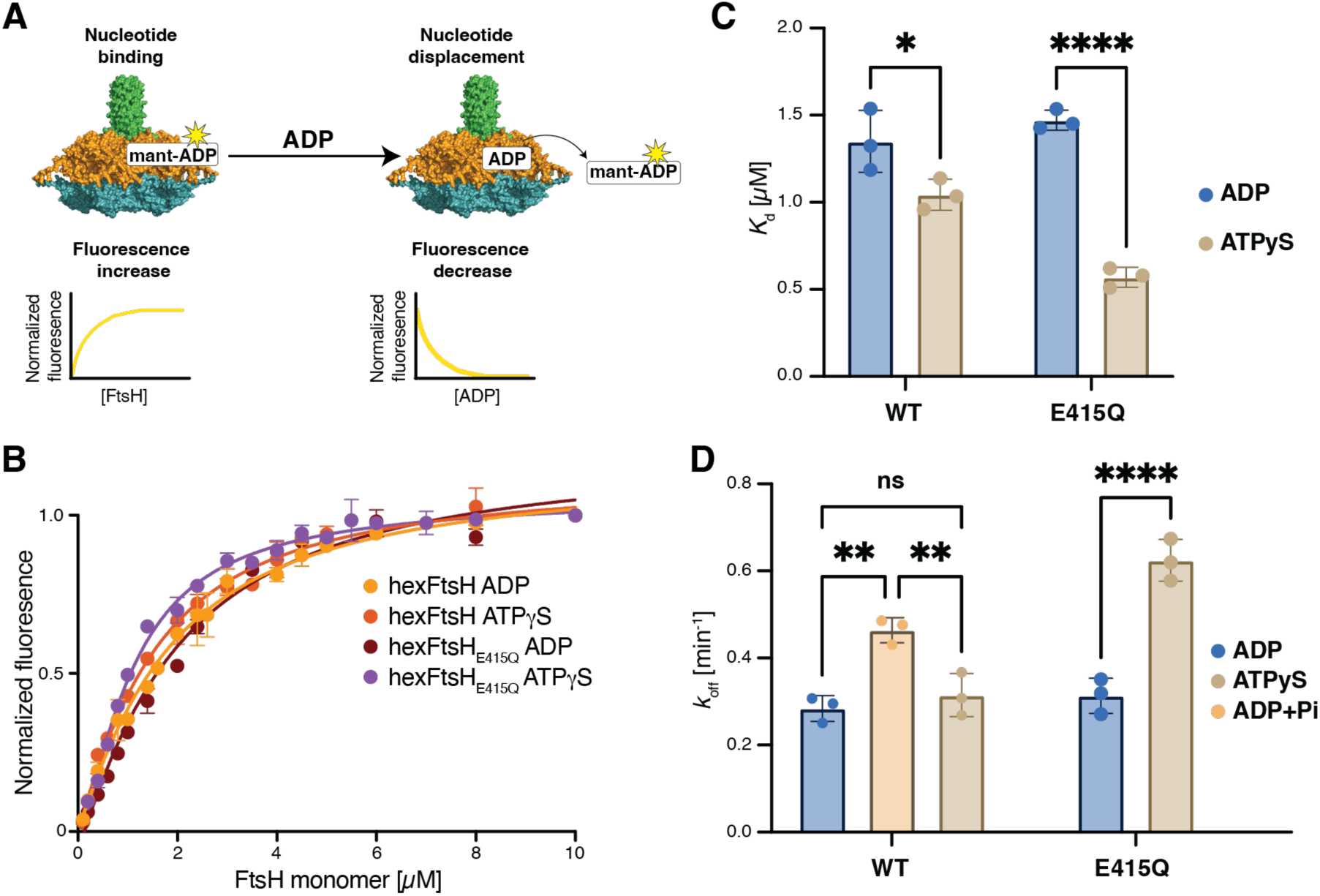
Assessing the nucleotide binding by fluorescence. **A**) Schematics of the employed fluorescence titration assays using fluorescently labelled mant-nucleotides to measure the dissociation constants (*K*_d_) as well as the off-rate (*k*_off_) by a displacement titration. **B**) Fluorescence binding curves obtained for mant-ADP and mant-ATPγS upon addition of hexFtsH or hexFtsH_E415Q_, respectively. **C**) Extracted dissociation constants, *K*_d_s, for hexFtsH and hexFtsH_E415Q_, respectively. Statistical significance was assessed *via* the one-way ANOVA test resulting in *p*= 0.016 for hexFtsH and *p*= <0.0001 for hexFtsH_E415Q_. **D**) Obtained nucleotide *k*_off_-rates derived from the mant-nucleotide displacement assay for hexFtsH and hexFtsH_E415Q_, respectively. Statistical significance was assessed *via* the one-way ANOVA test resulting in *p*= 0.0026, 0.6, 0.0064 for hexFtsH as well as *p*= <0.0001 for hexFtsH_E415Q_.

### Substrate induced stimulation of ATPase cycle reveals negative allosteric coupling to the catalytic site in wild-type FtsH

We next assessed the ATPase activity under steady state conditions for the hexFtsH constructs employing an NADH-coupled enzymatic assay used alongside an ATP regeneration system as described above for the NMR-assay in detail. We ran the experiment both in the absence and in the presence of the substrate β-casein (**Fig. 8**). For hexFtsH, the ATPase activity was determined to be 2 ± 0.06 min^-1^ hexamer^-1^ with an apparent *K*_m_ of 180 ± 17 μM. Upon substrate addition we observed an ∼10-fold increased ATPase activity to 26 ± 0.4 min^-1^ hexamer^-1^ with a 4–5-fold increased ATP affinity, *K*_m_ = 43 ± 3 μM, which is in line with the obtained trends in previous reports using alternative substrates (*80*). This substrate-induced upregulation of the ATPase rate accompanied by a ∼4-fold reduced *K*_m_ suggests β-casein-induced substrate dependent structural modulations within both FtsH-domains. Remarkably for the hexFtsH_E415Q_ mutant, the ATPase activity showed a notable increase compared to the wild-type hexFtsH variant, leading to an ATPase rate of 208 ± 3 min^-1^ hexamer^-1^ and a *K*_m_ = 116 ± 7 μM. This drastic increase further points to a direct coupling between the ATPase and the proteolytic site, with the latter apparently limiting the ATPase rate. Upon β-casein addition, the ATPase rate moderately decreased to 141 ± 4 min^-1^ hexamer^-1^ without a notable effect on the *K*_m_ with a value of 112 ± 11 μM, indicating that the ATPase regulation is directly dependent on the proteolytic state. Thus, in summary the E415Q-mutation impairs the interdomain allosteric communication as hexFtsH_E415Q_ has an abolished proteolytic activity and thus cannot degrade the substrate, whereas this lack of turnover capacity appears to be reflected by the observed moderate decrease of *k*_cat_. Nevertheless, it is still able to bind and hydrolyze ATP needed to unfold and translocate the substrate. The decrease of ATP turnover upon addition of β-casein can to some extent be expected as the hexFtsH_E415Q_ likely remains locked in a substrate engaged state since it cannot process and subsequently release it.

**Figure 8.**
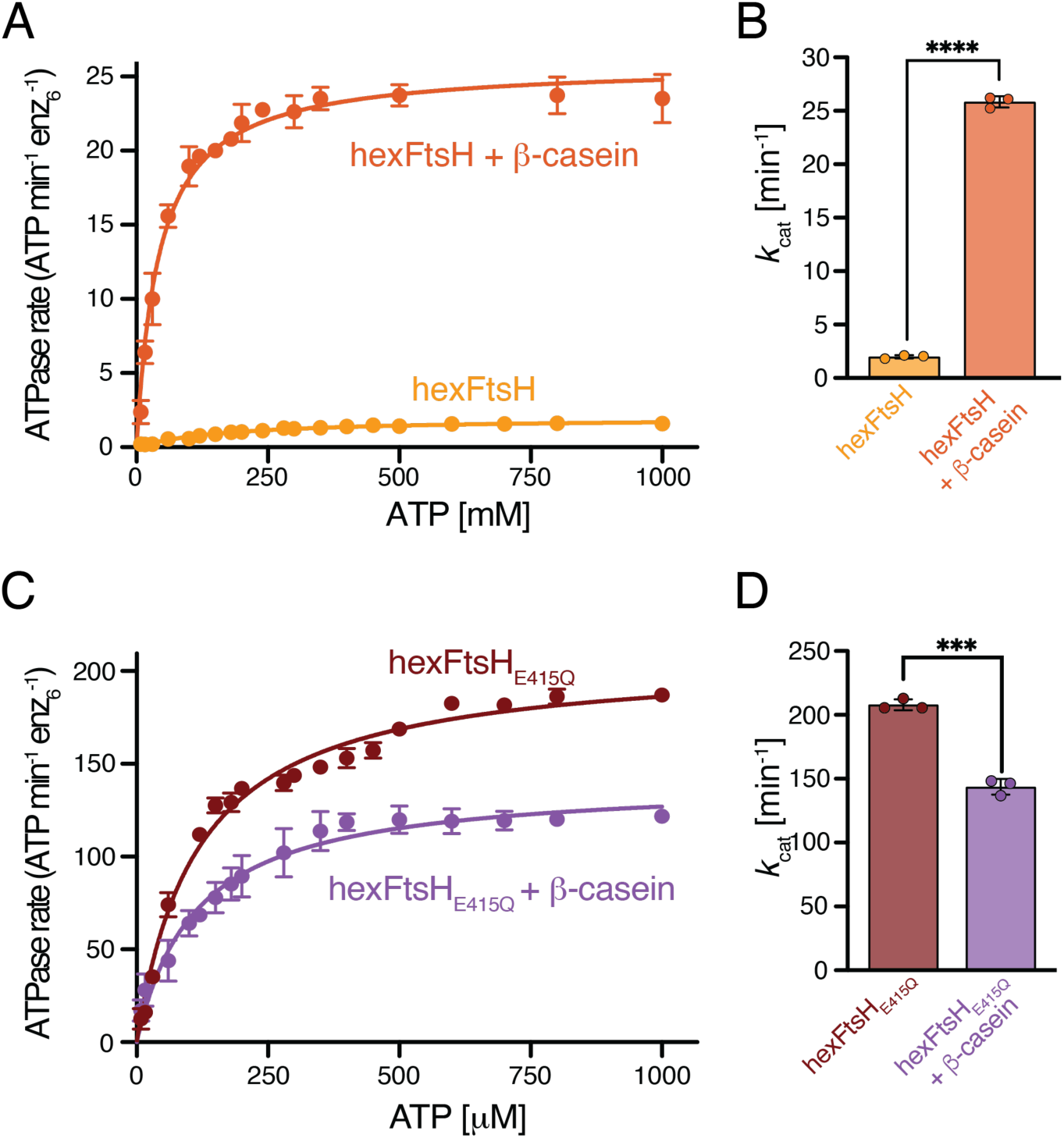
Stimulated ATPase activity. **A)** Fluorescence curves of the NADH coupled ATPase assay in the absence (yellow) and the presence (orange) of β-casein using hexFtsH. **B**) Obtained *k*_cat_ values with and without β-casein. Statistical significance was assessed *via* the one-way ANOVA test resulting in *p*< 0.0001. **C)** Fluorescence curves of an NADH coupled ATPase in the absence (dark-red) and the presence (purple) of β-casein using hexFtsH_E415Q_. **D**) Obtained *k*_cat_ values with and without β-casein. Statistical significance was assessed *via* the one-way ANOVA test resulting in *p*=0.0001. All experiments were performed as biological triplicates.

## Conclusion

Whereas the functional cycle of soluble AAA+ ATPases has been studied at detail the understanding of the underlying functional principles of membrane associated system remains sparse (*64*). Here, we elucidate by using advanced solution NMR spectroscopy approaches the intricate regulation of the FtsH ATPase domain at unprecedented level of detail. Our data revealed the functional interplay between the two sub-domains of this important functional domain and outline an allosteric feedback loop to the protease domain. Methyl NMR-spectroscopy enabled us to reveal crucial interactions facilitating the communication between different parts of the ATPase module based on altered dynamical adaptations.

Using a combination of *in-cyclo* NMR and fluorescence-based assays enabled us to delineate the details of the FtsH ATPase cycle. This approach resulted in an almost complete picture of kinetic rates enabling quantitative description of the functional cycle (**Fig. 9**). We obtained clear evidence that the ATP hydrolysis is not the rate limiting step of the enzymatic reaction. This was seen as, on the one hand our data clearly show that the catalytic cycle for the hexamer is faster than the ATP hydrolysis at individual sites, pointing to positive cooperativity. And on the other hand, the observed feedback regulation between the protease and the ATPase sites, which become unhinged for the hexFtsH_E415Q_ variant. Even though the 51 s for a full ATP consumption cycle might seem to be quite extensive, this value is comparable to a recent study on the human Hsp70 protein BIP, where the authoŕs obtained similar values (e.g. ∼40 s) and could show an acceleration of the cycle in the presence of nucleotide exchange factors and other co-chaperones (*82*).

**Figure 9.**
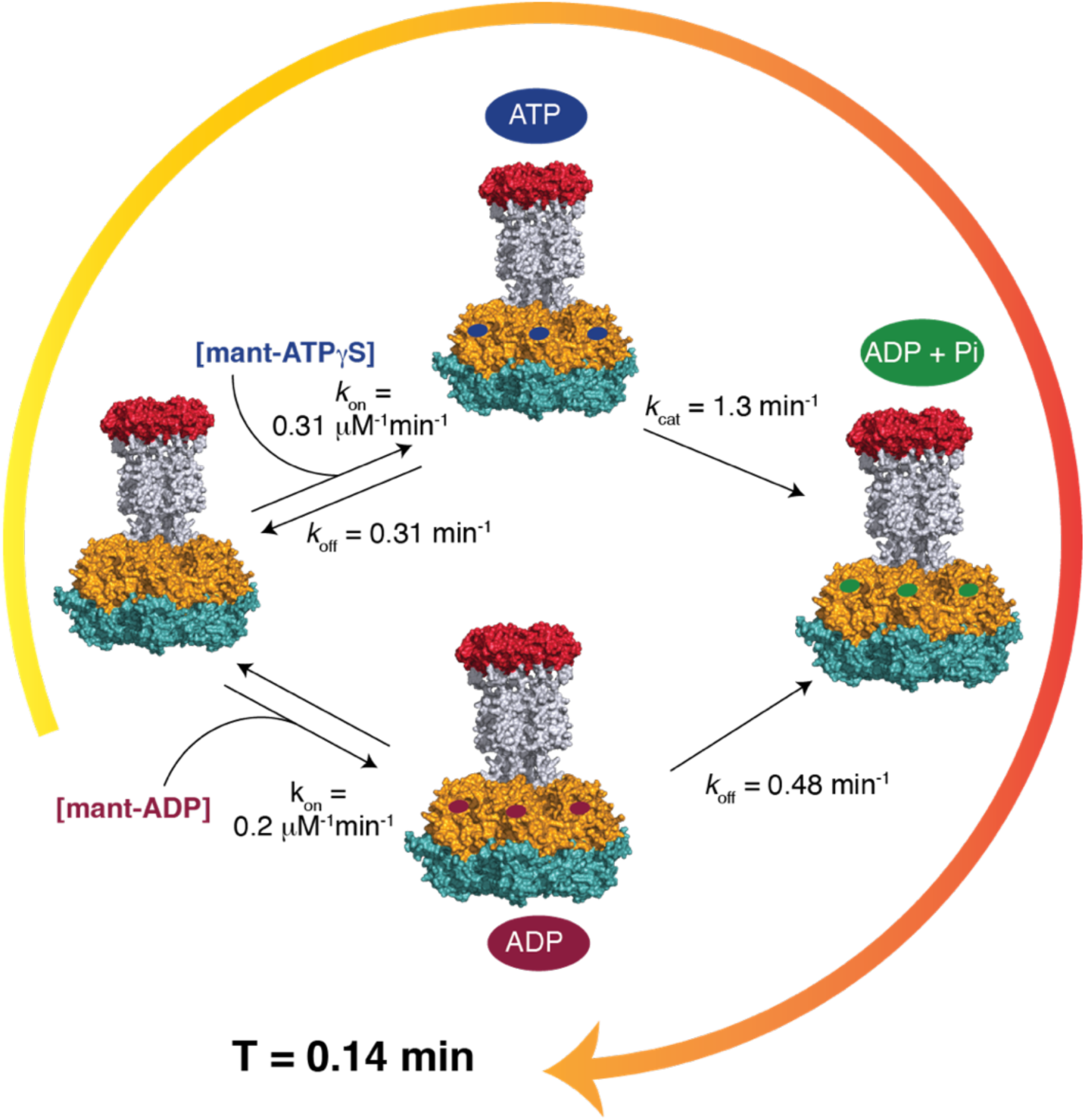
The four-step functional ATP-cycle of FtsH. The AAA+ protease FtsH cycles through four states during ATP consumption. The kinetic parameters were established by the different assays described herein. The non/slow hydrolysable ATP analogue, ATPγS is used to describe ATP binding.

As we obtained similar dissociation constants for mant-ADP and mant-ATPγS as well as comparable lifetimes, we hypothesized that the product release might be the rate limiting step. Indeed, we could observe in the presence of phosphate a faster release of ADP, thus a destabilization of the product state, supporting product-release being the rate limiting step. Previous studies suggested that the hydrolysis might be modulated by the so-called arginine finger of a neighbouring sub-unit, which is in line with our observation and indicates a role for this motif in the release of the products (*90*). In the context of other AAA+ ATPases, this observation suggests that the product-release is commonly the rate limiting step, likely due to a high kinetic barrier to phosphate dissociation (*65*, *91*).

## Acknowledgement

The Swedish NMR Centre of the University of Gothenburg, supported through the Swedish Research Council financed SwedNMR infrastructure consortium and the SciLifeLab Integrated Structural Biology (ISB) platform is gratefully acknowledged for spectrometer time. Research in the Burmann lab is supported by funding from the Swedish Research Council (Starting Grant 2016–04721; Consolidator Grant 2020–00466), the Swedish Cancer Foundation (2019-0415 and 2022–2490), and the Knut och Alice Wallenberg Foundation through a Wallenberg Academy Fellowship (2016.0163 and 2020.0300) as well as through the Wallenberg Centre for Molecular and Translational Medicine, University of Gothenburg Sweden. B.M.B. gratefully acknowledges an EMBO Young Investigator Fellowship.

## Data and materials availability

All data needed to evaluate the conclusions in the paper are present in the paper and/or the Supplementary Materials. The sequence-speciÉic NMR resonance assignments for the aFtsH were deposited in the BioMagResBank (www.bmrb.wisc.edu) under accession codes XXX. The NMR data used for methyl group analyis and relaxation analysis have been tabulated and are available alongsidethe ConSurf generated MSA (multiple sequence alignment) Éile on XX.

## References

1. M. S. Hipp, P. Kasturi, F. U. Hartl, The proteostasis network and its decline in ageing. Nat. Rev. Mol. Cell. Biol. 20, 421–435 (2019).

2. F. U. Hartl, A. Bracher, M. Hayer-Hartl, Molecular chaperones in protein folding and proteostasis. Nature 475, 324–332 (2011).

3. R. T. Sauer, T. A. Baker, AAA+ Proteases: ATP-Fueled Machines of Protein Destruction. Annu. Rev. Biochem. 80, 587–612 (2011).

4. A. O. Olivares, T. A. Baker, R. T. Sauer, Mechanistic insights into bacterial AAA+ proteases and protein-remodelling machines. Nat. Rev. Microbiol. 14, 33–44 (2016).

5. L. Bittner, J. Arends, F. Narberhaus, Mini review: ATP-dependent proteases in bacteria. Biopolymers. 105, 505–517 (2016).

6. K. Nyquist, A. Martin, Marching to the beat of the ring: polypeptide translocation by AAA+ proteases. Trends. Biochem. Sci. 39, 53–60 (2014).

7. K. Ito, Y. Akiyama, Cellular functions, mechanism of action, and regulation of FtsH protease. Annu. Rev. Microbiol. 59, 211–231 (2005).

8. L.-M. Bittner, J. Arends, F. Narberhaus, When, how and why? Regulated proteolysis by the essential FtsH protease in *Escherichia coli*. Biol. Chem. 398, 625–635 (2017).

9. W. Liu, M. Schoonen, T. Wang, S. McSweeney, Q. Liu, Cryo-EM structure of transmembrane AAA+ protease FtsH in the ADP state. *Commun*. Biol. 5, 257 (2022).

10. S. Langklotz, U. Baumann, F. Narberhaus, Structure and function of the bacterial AAA protease FtsH. Biochim. Biophys. Acta Mol. Cell Res. 1823, 40–48 (2012).

11. S. Krzywda, A. M. Brzozowski, C. Verma, K. Karata, T. Ogura, A. J. Wilkinson, The Crystal Structure of the AAA Domain of the ATP-Dependent Protease FtsH of Escherichia coli at 1.5 Å Resolution. Structure 10, 1073–1083 (2002).

12. T. Ogura, A. J. Wilkinson, AAA^+^ superfamily ATPases: common structure–diverse function. Genes Cells. 6, 575–597 (2001).

13. T. Langer, AAA proteases: cellular machines for degrading membrane proteins. Trends Biochem. Sci. 25, 247–251 (2000).

14. Z. Qiao, T. Yokoyama, X.-F. Yan, I. T. Beh, J. Shi, S. Basak, Y. Akiyama, Y.-G. Gao, Cryo-EM structure of the entire FtsH-HflKC AAA protease complex. Cell. Rep. 39, 110890 (2022).

15. M. K. Black, A. Kim, C. Y. Chen, M. M. Goncalves, S. Waseem, S. Vahidi, R. Huang, Characterization of Conformational Dynamics and Structural Plasticity of the Catalytic Domain of Human Mitochondrial YME1L Protease. Biochemistry. 65, 732–747 (2026).

16. M. Geiser, R. Cébe, D. Drewello, R. Schmitz, Integration of PCR Fragments at Any Specific Site within Cloning Vectors without the Use of Restriction Enzymes and DNA Ligase. BioTechniques 31, 88–92 (2001).

17. J. Mikolajczyk, M. Drag, M. Békés, J. T. Cao, Z. Ronai, G. S. Salvesen, Small Ubiquitin-related Modifier (SUMO)-specific Proteases. J. Biol. Chem. 282, 26217–26224 (2007).

18. M. Callon, B. M. Burmann, S. Hiller, Structural Mapping of a Chaperone–Substrate Interaction Surface. Angew. Chem. Int. Ed. 53, 5069–5072 (2014).

19. M. Salzmann, K. Pervushin, G. Wider, H. Senn, K. Wüthrich, TROSY in triple-resonance experiments: New perspectives for sequential NMR assignment of large proteins. Proc. Natl. Acad. Sci. U.S.A. 95, 13585–13590 (1998).

20. K. Pervushin, R. Riek, G. Wider, K. Wüthrich, Attenuated *T* _2_ relaxation by mutual cancellation of dipole–dipole coupling and chemical shift anisotropy indicates an avenue to NMR structures of very large biological macromolecules in solution. Proc. Natl. Acad. Sci. U.S.A. 94, 12366–12371 (1997).

21. M. Sattler, Heteronuclear multidimensional NMR experiments for the structure determination of proteins in solution employing pulsed field gradients. Prog. Nucl. Magn. Reson. Spectrosc. 34, 93–158 (1999).

22. P. Rossi, Y. Xia, N. Khanra, G. Veglia, C. G. Kalodimos, 15N and 13C- SOFAST-HMQC editing enhances 3D-NOESY sensitivity in highly deuterated, selectively [1H,13C]-labeled proteins. J. Biomol. NMR. 66, 259–271 (2016).

23. V. Jaravine, I. Ibraghimov, V. Yu Orekhov, Removal of a time barrier for high-resolution multidimensional NMR spectroscopy. Nat. Methods. 3, 605–607 (2006).

24. F. Delaglio, S. Grzesiek, Geerten W. Vuister, G. Zhu, J. Pfeifer, A. Bax, NMRPipe: A multidimensional spectral processing system based on UNIX pipes. J. Biomol. NMR. 6 (1995).

25. R. L. J. Keller, “Optimizing the process of nuclear magnetic resonance spectrum analysis and computer aided resonance assignment,” thesis, ETH Zurich (2005).

26. J. T. Nielsen, F. A. A. Mulder, POTENCI: prediction of temperature, neighbor and pH-corrected chemical shifts for intrinsically disordered proteins. J. Biomol. NMR. 70, 141–165 (2018).

27. B. M. Burmann, C. Wang, S. Hiller, Conformation and dynamics of the periplasmic membrane-protein–chaperone complexes OmpX–Skp and tOmpA–Skp. Nat. Struct. Mol. Biol. 20, 1265–1272 (2013).

28. L. Morgado, B. M. Burmann, T. Sharpe, A. Mazur, S. Hiller, The dynamic dimer structure of the chaperone Trigger Factor. Nat. Commun. 8, 1992 (2017).

29. G. Wider, L. Dreier, Measuring Protein Concentrations by NMR Spectroscopy. J. Am. Chem. Soc. 128, 2571–2576 (2006).

30. J. Lidman, Y. Sallova, I. Matečko-Burmann, B. M. Burmann, Structure and dynamics of the mitochondrial DNA-compaction factor Abf2 from S. cerevisiae. J. Struct. Biol. 215, 108008 (2023).

31. K. Takeuchi, Y. Tokunaga, M. Imai, H. Takahashi, I. Shimada, Dynamic multidrug recognition by multidrug transcriptional repressor LmrR. Sci. Rep. 4, 6922 (2014).

32. G. L. Butterfoss, E. F. DeRose, S. A. Gabel, L. Perera, J. M. Krahn, G. A. Mueller, X. Zheng, R. E. London, Conformational dependence of 13C shielding and coupling constants for methionine methyl groups. J. Biomol. NMR. 48, 31–47 (2010).

33. D. F. Hansen, P. Neudecker, L. E. Kay, Determination of Isoleucine Side-Chain Conformations in Ground and Excited States of Proteins from Chemical Shifts. J. Am. Chem. Soc. 132, 7589–7591 (2010).

34. N.-A. Lakomek, J. Ying, A. Bax, Measurement of 15N relaxation rates in perdeuterated proteins by TROSY-based methods. J. Biomol. NMR. 53, 209–221 (2012).

35. M. Niklasson, R. Otten, A. Ahlner, C. Andresen, J. Schlagnitweit, K. Petzold, P. Lundström, Comprehensive analysis of NMR data using advanced line shape fitting. J. Biomol. NMR. 69, 93–99 (2017).

36. P. Dosset, J.-C. Hus, M. Blackledge, D. Marion, Efficient analysis of macromolecular rotational diffusion from heteronuclear relaxation data. J. Biomol. NMR. 16, 23–28 (2000).

37. M. W. Maciejewski, A. D. Schuyler, M. R. Gryk, I. I. Moraru, P. R. Romero, E. L. Ulrich, H. R. Eghbalnia, M. Livny, F. Delaglio, J. C. Hoch, NMRbox: A Resource for Biomolecular NMR Computation. Biophys. J. 112, 1529–1534 (2017).

38. R. Cole, J. P. Loria, FAST-Modelfree: A program for rapid automated analysis of solution NMR spin-relaxation data. J. Biomol. NMR. 26, 203–213 (2003).

39. J. García De La Torre, M. L. Huertas, B. Carrasco, Calculation of Hydrodynamic Properties of Globular Proteins from Their Atomic-Level Structure. Biophys. J. 78, 719–730 (2000).

40. A. A. Smith, M. Ernst, B. H. Meier, Optimized “detectors” for dynamics analysis in solid-state NMR. J. Chem. Phys. 148, 045104 (2018).

41. A. A. Smith, M. Ernst, B. H. Meier, F. Ferrage, Reducing bias in the analysis of solution-state NMR data with dynamics detectors. J. Chem. Phys. 151, 034102 (2019).

42. V. Tugarinov, L. E. Kay, An Isotope Labeling Strategy for Methyl TROSY Spectroscopy. J. Biomol. NMR. 28, 165–172 (2004).

43. K. Weinhäupl, C. Lindau, A. Hessel, Y. Wang, C. Schütze, T. Jores, L. Melchionda, B. Schönfisch, H. Kalbacher, B. Bersch, D. Rapaport, M. Brennich, K. Lindorff-Larsen, N. Wiedemann, P. Schanda, Structural Basis of Membrane Protein Chaperoning through the Mitochondrial Intermembrane Space. Cell. 175, 1365–1379.e25 (2018).

44. H. Sun, L. E. Kay, V. Tugarinov, An Optimized Relaxation-Based Coherence Transfer NMR Experiment for the Measurement of Side-Chain Order in Methyl-Protonated, Highly Deuterated Proteins. J. Phys. Chem. B. 115, 14878–14884 (2011).

45. D. M. Korzhnev, K. Kloiber, V. Kanelis, V. Tugarinov, L. E. Kay, Probing Slow Dynamics in High Molecular Weight Proteins by Methyl-TROSY NMR Spectroscopy: Application to a 723-Residue Enzyme. J. Am. Chem. Soc. 126, 3964–3973 (2004).

46. C. A. Schneider, W. S. Rasband, K. W. Eliceiri, NIH Image to ImageJ: 25 years of image analysis. Nat. Methods. 9, 671–675 (2012).

47. E. Agustoni, R. D. Teixeira, M. Huber, S. Flister, S. Hiller, T. Schirmer, Acquisition of enzymatic progress curves in real time by quenching-free ion exchange chromatography. Anal. Biochem. 639, 114523 (2022).

48. C. Ma, C. Wang, D. Luo, L. Yan, W. Yang, N. Li, N. Gao, Structural insights into the membrane microdomain organization by SPFH family proteins. Cell. Res. 32, 176–189 (2022).

49. K. Wüthrich, NMR assignments as a basis for structural characterization of denatured states of globular proteins. Curr. Opin. Struct. Biol. 4, 93–99 (1994).

50. J. Abramson, J. Adler, J. Dunger, R. Evans, T. Green, A. Pritzel, O. Ronneberger, L. Willmore, A. J. Ballard, J. Bambrick, S. W. Bodenstein, D. A. Evans, C.-C. Hung, M. O’Neill, D. Reiman, K. Tunyasuvunakool, Z. Wu, A. Žemgulytė, E. Arvaniti, C. Beattie, O. Bertolli, A. Bridgland, A. Cherepanov, M. Congreve, A. I. Cowen-Rivers, A. Cowie, M. Figurnov, F. B. Fuchs, H. Gladman, R. Jain, Y. A. Khan, C. M. R. Low, K. Perlin, A. Potapenko, P. Savy, S. Singh, A. Stecula, A. Thillaisundaram, C. Tong, S. Yakneen, E. D. Zhong, M. Zielinski, A. Žídek, V. Bapst, P. Kohli, M. Jaderberg, D. Hassabis, J. M. Jumper, Accurate structure prediction of biomolecular interactions with AlphaFold 3. Nature 630, 493–500 (2024).

51. A. G. Palmer, C. D. Kroenke, J. Patrick Loria, “Nuclear Magnetic Resonance Methods for Quantifying Microsecond-to-Millisecond Motions in Biological Macromolecules” in Methods in Enzymology (Elsevier, 2001)vol. 339, pp. 204–238.

52. F. Ferrage, A. Piserchio, D. Cowburn, R. Ghose, On the measurement of 15N–{1H} nuclear Overhauser effects. J. Magn. Reson. 192, 302–313 (2008).

53. P. Rossi, G. V. T. Swapna, Y. J. Huang, J. M. Aramini, C. Anklin, K. Conover, K. Hamilton, R. Xiao, T. B. Acton, A. Ertekin, J. K. Everett, G. T. Montelione, A microscale protein NMR sample screening pipeline. J. Biomol. NMR. 46, 11–22 (2010).

54. C. Bieniossek, T. Schalch, M. Bumann, M. Meister, R. Meier, U. Baumann, The molecular architecture of the metalloprotease FtsH. Proc. Natl. Acad. Sci. U.S.A. 103, 3066–3071 (2006).

55. C. Bieniossek, B. Niederhauser, U. M. Baumann, The crystal structure of *apo* -FtsH reveals domain movements necessary for substrate unfolding and translocation. Proc. Natl. Acad. Sci. U.S.A. 106, 21579–21584 (2009).

56. G. M. Clore, A. Szabo, A. Bax, L. E. Kay, P. C. Driscoll, A. M. Gronenborn, Deviations from the simple two-parameter model-free approach to the interpretation of nitrogen-15 nuclear magnetic relaxation of proteins. J. Am. Chem. Soc. 112, 4989–4991 (1990).

57. G. Lipari, A. Szabo, Model-free approach to the interpretation of nuclear magnetic resonance relaxation in macromolecules. 1. Theory and range of validity. J. Am. Chem. Soc. 104, 4546–4559 (1982).

58. G. Lipari, A. Szabo, Model-free approach to the interpretation of nuclear magnetic resonance relaxation in macromolecules. 2. Analysis of experimental results. J. Am. Chem. Soc. 104, 4559–4570 (1982).

59. A. G. Palmer, F. Massi, Characterization of the Dynamics of Biomacromolecules Using Rotating-Frame Spin Relaxation NMR Spectroscopy. Chem. Rev. 106, 1700–1719 (2006).

60. H. K. Wayment-Steele, G. El Nesr, R. Hettiarachchi, A. M. Ojoawo, H. Kariyawasam, S. Ovchinnikov, D. Kern, Learning millisecond protein dynamics from what is missing in NMR spectra. Biophysics [Preprint] (2025). 10.1101/2025.03.19.642801.

61. R. G. Parra, N. P. Schafer, L. G. Radusky, M.-Y. Tsai, A. B. Guzovsky, P. G. Wolynes, D. U. Ferreiro, Protein Frustratometer 2: a tool to localize energetic frustration in protein molecules, now with electrostatics. Nucleic. Acids. Res. 44, W356–W360 (2016).

62. M. Landau, I. Mayrose, Y. Rosenberg, F. Glaser, E. Martz, T. Pupko, N. Ben-Tal, ConSurf 2005: the projection of evolutionary conservation scores of residues on protein structures. Nucleic. Acids. Res. 33, W299–W302 (2005).

63. H. Ashkenazy, E. Erez, E. Martz, T. Pupko, N. Ben-Tal, ConSurf 2010: calculating evolutionary conservation in sequence and structure of proteins and nucleic acids. Nucleic. Acids. Res. 38, W529–W533 (2010).

64. C. Puchades, C. R. Sandate, G. C. Lander, The molecular principles governing the activity and functional diversity of AAA+ proteins. Nat. Rev. Mol. Cell. Biol. 21, 43–58 (2020).

65. C. Puchades, A. J. Rampello, M. Shin, C. J. Giuliano, R. L. Wiseman, S. E. Glynn, G. C. Lander, Structure of the mitochondrial inner membrane AAA+ protease YME1 gives insight into substrate processing. Science 358, eaao0464 (2017).

66. C. Puchades, B. Ding, A. Song, R. L. Wiseman, G. C. Lander, S. E. Glynn, Unique Structural Features of the Mitochondrial AAA+ Protease AFG3L2 Reveal the Molecular Basis for Activity in Health and Disease. Mol. Cell. 75, 1073–1085.e6 (2019).

67. E. E. Aspholm, J. Lidman, B. M. Burmann, Structural basis of substrate recognition and allosteric activation of the proapoptotic mitochondrial HtrA2 protease. Nat. Commun. 15, 4592 (2024).

68. R. Rosenzweig, L. E. Kay, Bringing Dynamic Molecular Machines into Focus by Methyl-TROSY NMR. Annu. Rev. Biochem. 83, 291–315 (2014).

69. D. Šulskis, J. Thoma, B. M. Burmann, Structural basis of DegP protease temperature-dependent activation. Sci. Adv. 7, eabj1816 (2021).

70. A. B. Kleist, S. Jenjak, A. Sente, L. J. Laskowski, M. Szpakowska, M. M. Calkins, E. I. Anderson, L. M. McNally, R. Heukers, V. Bobkov, F. C. Peterson, M. A. Thomas, A. Chevigné, M. J. Smit, J. D. McCorvy, M. M. Babu, B. F. Volkman, Conformational selection guides β-arrestin recruitment at a biased G protein–coupled receptor. Science 377, 222–228 (2022).

71. C. C. Valley, A. Cembran, J. D. Perlmutter, A. K. Lewis, N. P. Labello, J. Gao, J. N. Sachs, The Methionine-aromatic Motif Plays a Unique Role in Stabilizing Protein Structure. J. Biol. Chem. 287, 34979–34991 (2012).

72. A. K. Lewis, K. M. Dunleavy, T. L. Senkow, C. Her, B. T. Horn, M. A. Jersett, R. Mahling, M. R. McCarthy, G. T. Perell, C. C. Valley, C. B. Karim, J. Gao, W. C. K. Pomerantz, D. D. Thomas, A. Cembran, A. Hinderliter, J. N. Sachs, Oxidation increases the strength of the methionine-aromatic interaction. Nat. Chem. Biol. 12, 860–866 (2016).

73. R. E. London, B. D. Wingad, G. A. Mueller, Dependence of Amino Acid Side Chain^13^ C Shifts on Dihedral Angle: Application to Conformational Analysis. J. Am. Chem. Soc. 130, 11097–11105 (2008).

74. F. Gaudreault, M. Chartier, R. Najmanovich, Side-chain rotamer changes upon ligand binding: common, crucial, correlate with entropy and rearrange hydrogen bonding. Bioinformatics 28, i423–i430 (2012).

75. K. Teilum, M. B. A. Kunze, S. Erlendsson, B. B. Kragelund, (S)Pinning down protein interactions by NMR. Protein Sci. 26, 436–451 (2017).

76. M. Vostrukhina, A. Popov, E. Brunstein, M. A. Lanz, R. Baumgartner, C. Bieniossek, M. Schacherl, U. Baumann, The structure of *Aquifex aeolicus* FtsH in the ADP-bound state reveals a *C* _2_ -symmetric hexamer. Acta. Crystallogr. D. Biol. Crystallogr. 71, 1307–1318 (2015).

77. H. Shi, A. J. Rampello, S. E. Glynn, Engineered AAA+ proteases reveal principles of proteolysis at the mitochondrial inner membrane. Nat. Commun. 7, 13301 (2016).

78. K. Karata, T. Inagawa, A. J. Wilkinson, T. Tatsuta, T. Ogura, Dissecting the Role of a Conserved Motif (the Second Region of Homology) in the AAA Family of ATPases. J. Biol. Chem. 274, 26225–26232 (1999).

79. J. P. Morehouse, T. A. Baker, R. T. Sauer, FTSH degrades dihydrofolate reductase by recognizing a partially folded species. Protein. Sci. 31, e4410 (2022).

80. R. C. Bruckner, P. L. Gunyuzlu, R. L. Stein, Coupled Kinetics of ATP and Peptide Hydrolysis by *Escherichia coli* FtsH Protease. Biochemistry 42, 10843–10852 (2003).

81. C. Herman, S. Prakash, C. Z. Lu, A. Matouschek, C. A. Gross, Lack of a Robust Unfoldase Activity Confers a Unique Level of Substrate Specificity to the Universal AAA Protease FtsH. Mol. Cell. 11, 659–669 (2003).

82. G. Mas, S. Hiller, Mechanism of ATP hydrolysis in the Hsp70 BiP nucleotide-binding domain. Nat. Commun. 16, 5086 (2025).

83. L. Rohland, R. Kityk, L. Smalinskaitė, M. P. Mayer, Conformational dynamics of the Hsp70 chaperone throughout key steps of its ATPase cycle. Proc. Natl. Acad. Sci. U.S.A. 119, e2123238119 (2022).

84. R. Zananiri, S. Mangapuram Venkata, V. Gaydar, D. Yahalom, O. Malik, S. Rudnizky, O. Kleifeld, A. Kaplan, A. Henn, Auxiliary ATP binding sites support DNA unwinding by RecBCD. Nat. Commun. 13, 1806 (2022).

85. S. A. Mahmoud, B. Aldikacti, P. Chien, ATP hydrolysis tunes specificity of a AAA+ protease. Cell. Rep. 40, 111405 (2022).

86. H. Yaginuma, S. Kawai, K. V. Tabata, K. Tomiyama, A. Kakizuka, T. Komatsuzaki, H. Noji, H. Imamura, Diversity in ATP concentrations in a single bacterial cell population revealed by quantitative single-cell imaging. Sci. Rep. 4, 6522 (2014).

87. K. R. Albe, M. H. Butler, B. E. Wright, Cellular concentrations of enzymes and their substrates. J. Theor. Biol. 143, 163–195 (1990).

88. R. G. Shulman, T. R. Brown, K. Ugurbil, S. Ogawa, S. M. Cohen, J. A. Den Hollander, Cellular Applications of^31^ P and^13^ C Nuclear Magnetic Resonance. Science 205, 160–166 (1979).

89. N. Amin, A. Peterkofsky, A Dual Mechanism for Regulating cAMP Levels in Escherichia coli. J. Biol. Chem. 270, 11803–11805 (1995).

90. T. Ogura, S. W. Whiteheart, A. J. Wilkinson, Conserved arginine residues implicated in ATP hydrolysis, nucleotide-sensing, and inter-subunit interactions in AAA and AAA+ ATPases. J. Struct. Biol. 146, 106–112 (2004).

91. M. Shein, M. Hitzenberger, T. C. Cheng, S. R. Rout, K. D. Leitl, Y. Sato, M. Zacharias, E. Sakata, A. K. Schütz, Characterizing ATP processing by the AAA+ protein p97 at the atomic level. Nat. Chem. 16, 363–372 (2024).

