## Supplementary Material for "Functional details of the integral membrane metallo-protease FtsH revealed by solution NMR spectroscopy"

[*]Correspondence should be addressed to BMB:

**Supplementary Tables**

**Supplementary Table S1**: Details of the expression plasmids used in this study.

| **Plasmid** | **Tag** |
|---|---|
| pET28b-cFtsH | N-terminal $His_6$ |
| pET28b-pFtsH | N-terminal $His_6$-SUMO |
| pET28b-aFtsH | N-terminal $His_6$-SUMO |
| pET28b-hexFtsH | N-terminal $His_6$ |
| pET28b-hexFtsH$_{E415Q}$ | N-terminal $His_6$ |
| pET28b-hexFtsH$_{K201N}$ | N-terminal $His_6$ |

**Supplementary Table S2:** Primers used in this study to create the different FtsH constructs.

| **Plasmid** | **Primers** |
|---|---|
| pET28b-pFtsH | 5'-GCATCTGGAACAGATTGGCGGTGAAGCGCAGAAAGAATCG-3'<br>5'-GGTGGTGGTGCTCGAGTCACTTGTCGCCTAACTGCTCTGAC-3' |
| pET28b-aFtsH | 5'-CATCTGGAACAGATTGGCGGTATGGCGCGCATGCTGACGGAAG-3'<br>5'-GGTGGTGGTGGTGCTCGAGTCAACGTTCCGCTAGCATCATG-3' |
| pET28b-hexFtsH | 5'-GGTTCCTACTTCCAAAGCAACGCTGCGCGCATGCTGACGGAAG-3'<br>5'-GTGGTGGTGGTGGTGCTCGAGTTACTTGTCGCCTAACTGCTCTGACATGG-3' |
| pET28b-hexFtsH $_{E415Q}$ | 5'-CTTACCACCAGGCGGGTCATGCGATTATC-3'<br>5'-CCCGCCTGGTGGTAAGCCGTCGATTC-3' |
| pET28b-hexFtsH $_{K201N}$ | 5'-CCGGTAACACGCTGCTGGCGAAAGCG-3'<br>5'-CAGCGTGTTACCGGTACCCGGAGGACCG-3' |

**Supplementary Table S3:** Buffers used for purification and functional assays as indicated in the methods section.

| **Buffer** | **Composition** |
|---|---|
| A | 20 mM Tris, 500 mM NaCl, 1 mM DTT and 5 mM Imidazole, pH 8 |
| B | 50 mM Tris, 300 mM NaCl, 1 mM DTT, 10% glycerol, pH8 |
| C | 50 mM potassium phosphate, 300 mM KCl, pH 7.4 |
| D | 20 mM Tris, 150 mM NaCl, 5 mM $MgCl_2$, 1 mM DTT, 10% glycerol, pH 7.8 |
| E | 25 mM HEPES, 150 mM KCl, 5 mM $MgCl_2$, 25 mM $ZnCl_2$, 1 mM DTT, pH 7.4 |

**Supplementary Table S4:** Calculated molecular weights and estimated errors (S.D.) as determined by SEC-MALS for cFtsH, aFtsH, and pFtsH at room temperature in buffer C.

| **Construct** | **Mw [kDa]** |
|---|---|
| cFtsH | $56.3 \pm 0.56$ |
| pFtsH | $24.8 \pm 0.10$ |
| aFtsH ($1^{st}$ peak) | $55.1 \pm 0.72$ |
| aFtsH ($2^{nd}$ peak) | $26.3 \pm 0.13$ |

### Supplementary Figures

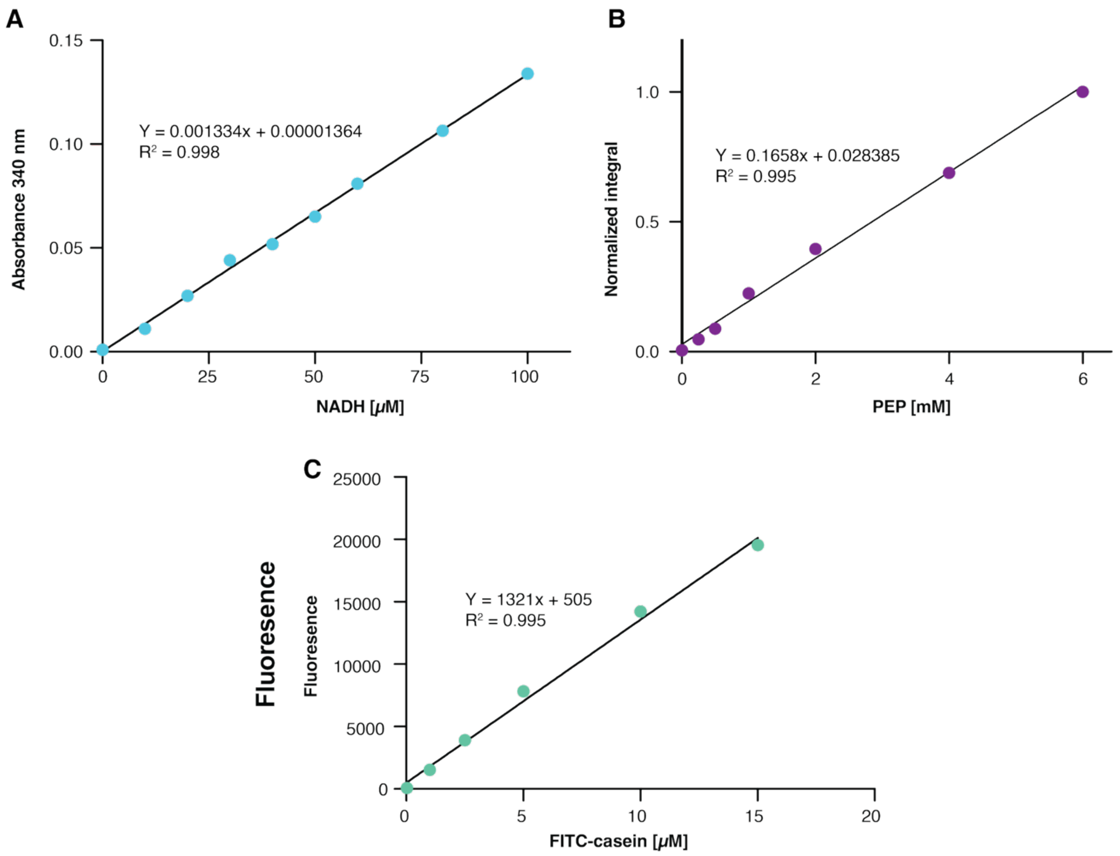


**Supplementary Figure S1. A)** NADH standard curve used to quantify ATP hydrolysis in the NADH-coupled steady-state ATP assay. Absorbance at 340 nm was plotted as a function of NADH concentration, and a linear fit was applied (equation depicted in the figure). The standard curve was used to convert absorbance values to NADH concentrations. **B)** Standard curve for phosphoenolpyruvate (PEP) used in the *in-cyclo* NMR based ATP regeneration assay. Normalized integral of PEP peak was plotted against concentration (equation for linear fit is indicated), the fit was used to convert integrals of the pyruvate peaks to concentrations to calculate the ATP turnover. **C**) FITC-casein standard curve. The standard curve was produced by trypsin-mediated degradation of different concentrations (15 μM, 10 μM, 5 μM, 2.5 μM, 1 μM, 0 μM) of FITC-casein. The standard curve was used to convert RFU to molar quantities for kinetic studies of hexFtsH.

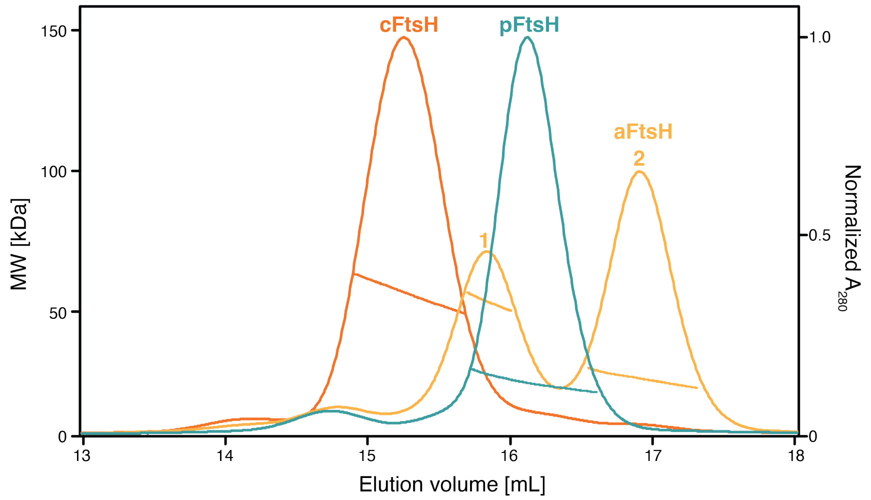


**Supplementary Figure S2.** SEC elution profiles of cFtsH (orange) pFtsH (blue) and aFtsH (yellow) plotted as MALS apparent molecular mass (left axis) and normalized absorbance ($A_{280}$) (right axis). Estimated molecular weights are reported in **Supplementary Table 4.**

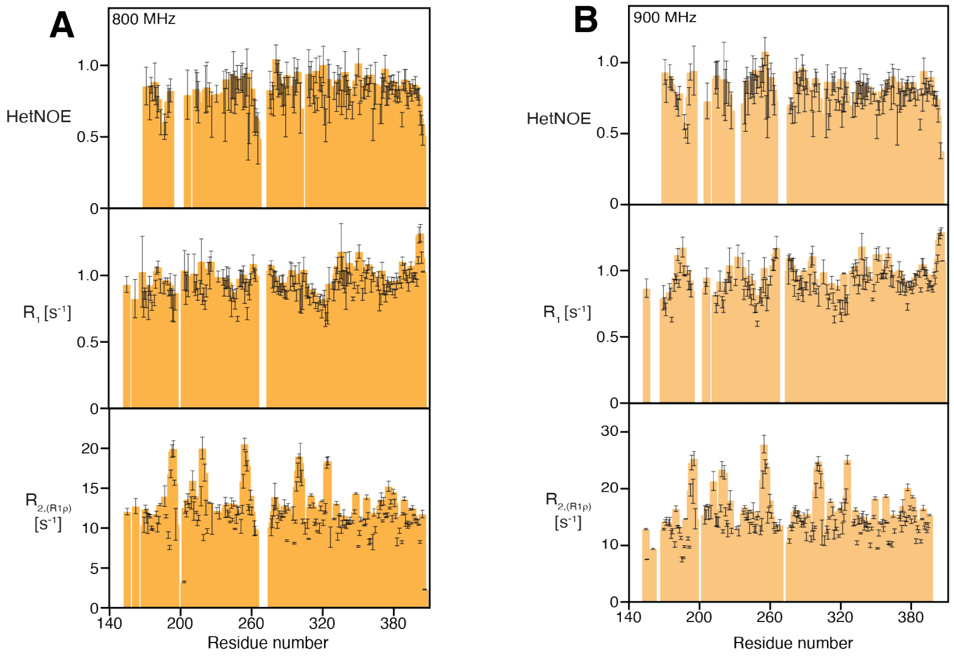


**Supplementary Figure S3.** Backbone dynamics plotted against the residue number. The pico- to nanosecond timescale probed by hetNOE (top) and longitudinal $R_1$ relaxation measurements (middle), alongside the micro- to millisecond dynamics probed by transcersal $R_{2(R1\rho)}$ measurements of **A)** aFtsH recorded at 18.8 T (800MHz). **B)** aFtsH recorded at 21.1 T (900MHz)..

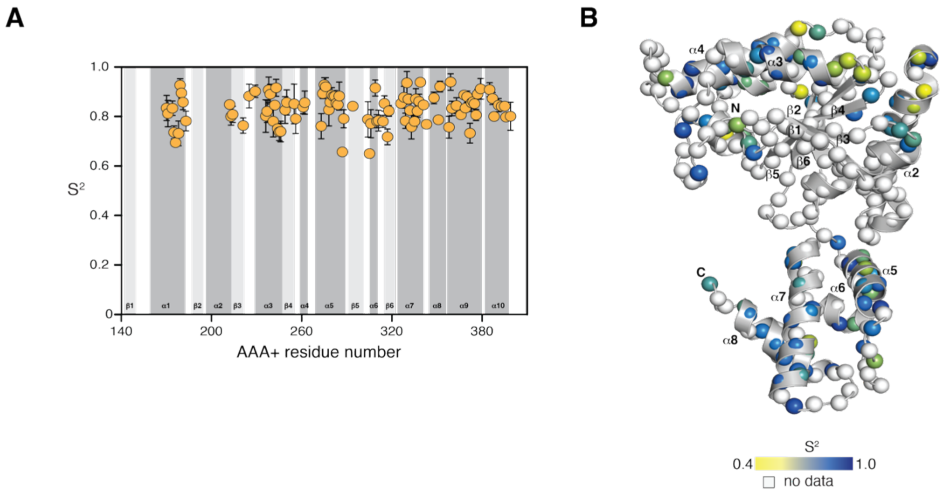


**Supplementary Figure S4. A)** The generalized order parameter $S^2$ reporting on pico- to nanosecond motions. **B)** $S^2$ values plotted onto the crystal structure of the AAA+ domain (PDB-ID: 1LV7). The amide moieties are shown as spheres and $S^2$ is indicated by the yellow-to-blue gradient.

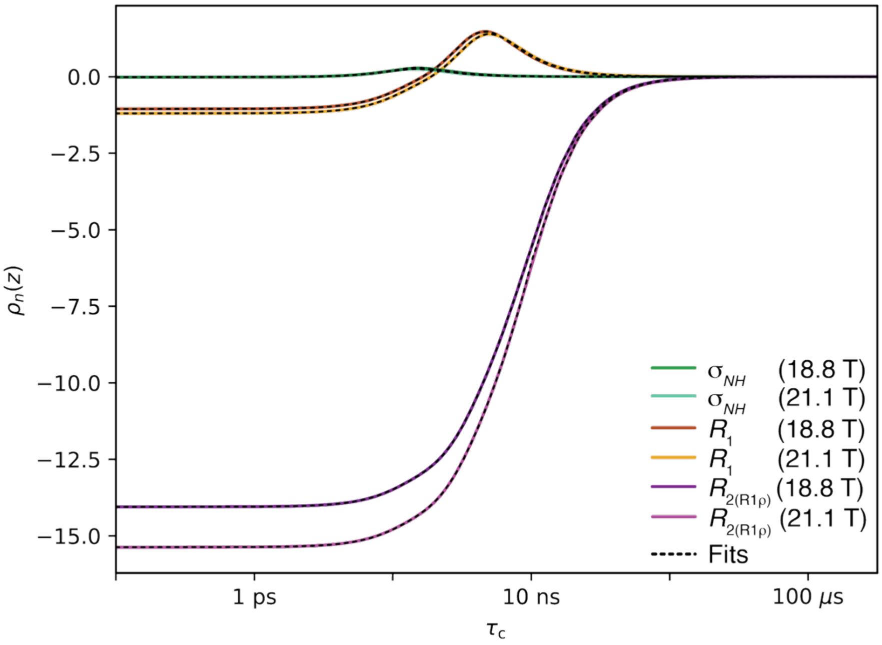


**Supplementary Figure S5.** Plotting of the responses ($\rho_n$(z)) of the measured relaxation parameters ($R_1$, $R_{2(R1\rho)}$, and $\sigma_{NH}$) measured at 18.8 T (800 MHz) and 21.1 T (900 MHz), respectively, plotted against the probed correlation times. Solid lines represent the experimental input data and the back-calculated data using 4 detectors (dotted lines). The amide cross correlation data, $\sigma_{NH}$, was determined through the following relationship: $\sigma_{NH}$ = (hetNOE – 1) • $R_1$ ($\gamma_N/\gamma_H$), with $\gamma_N/\gamma_H$ the ratio of the gyromagnetic ratio (-9.87), and hetNOE as well as $R_1$ the measured relaxation rates.

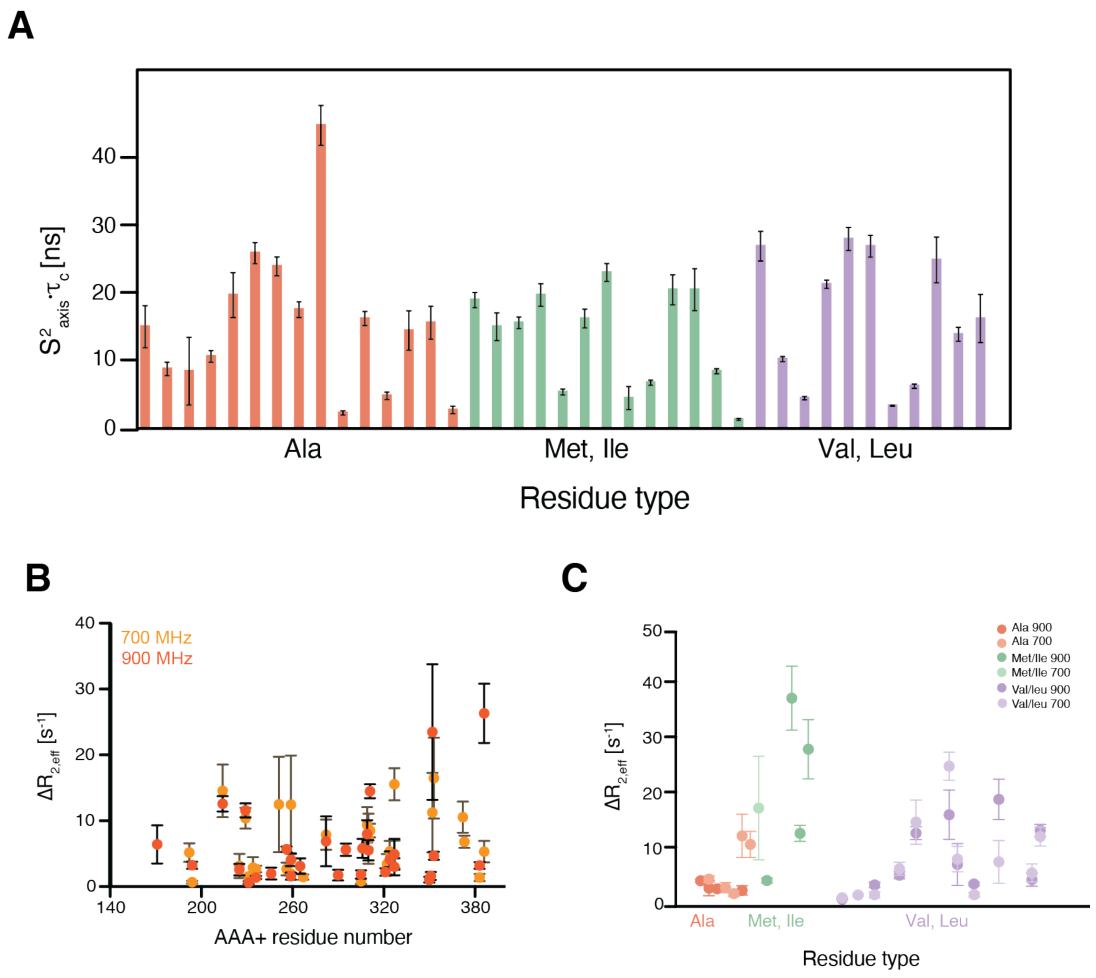


**Supplementary Figure S6. A)** Local methyl group dynamics of unassigned residues grouped by residue type, on the pico- to nanosecond timescale probed by methyl single quantum (SQ) and triple quantum (TQ) relaxation experiments showing the product of the local order parameter and the overall tumbling constant, $S^2_{axis} \bullet \tau_c$. **B)** $\Delta R_{2,eff}$ values for assigned methyl groups of the AAA+ domain in cFtsH recorded at 16.4 T (700 MHz) (yellow) and 21.1 T (900 MHz) (orange), obtained from the difference of the $R_{2,eff}$ at the lowest and the highest CPMG frequency. **C)** $\Delta R_{2,eff}$ values for unassigned methyl groups of cFtsH, grouped by residue type, recorded at 16.4 T (700 MHz) and 21.1 T (900 MHz), obtained from the difference of the $R_{2,eff}$ at the lowest and the highest CPMG frequency.

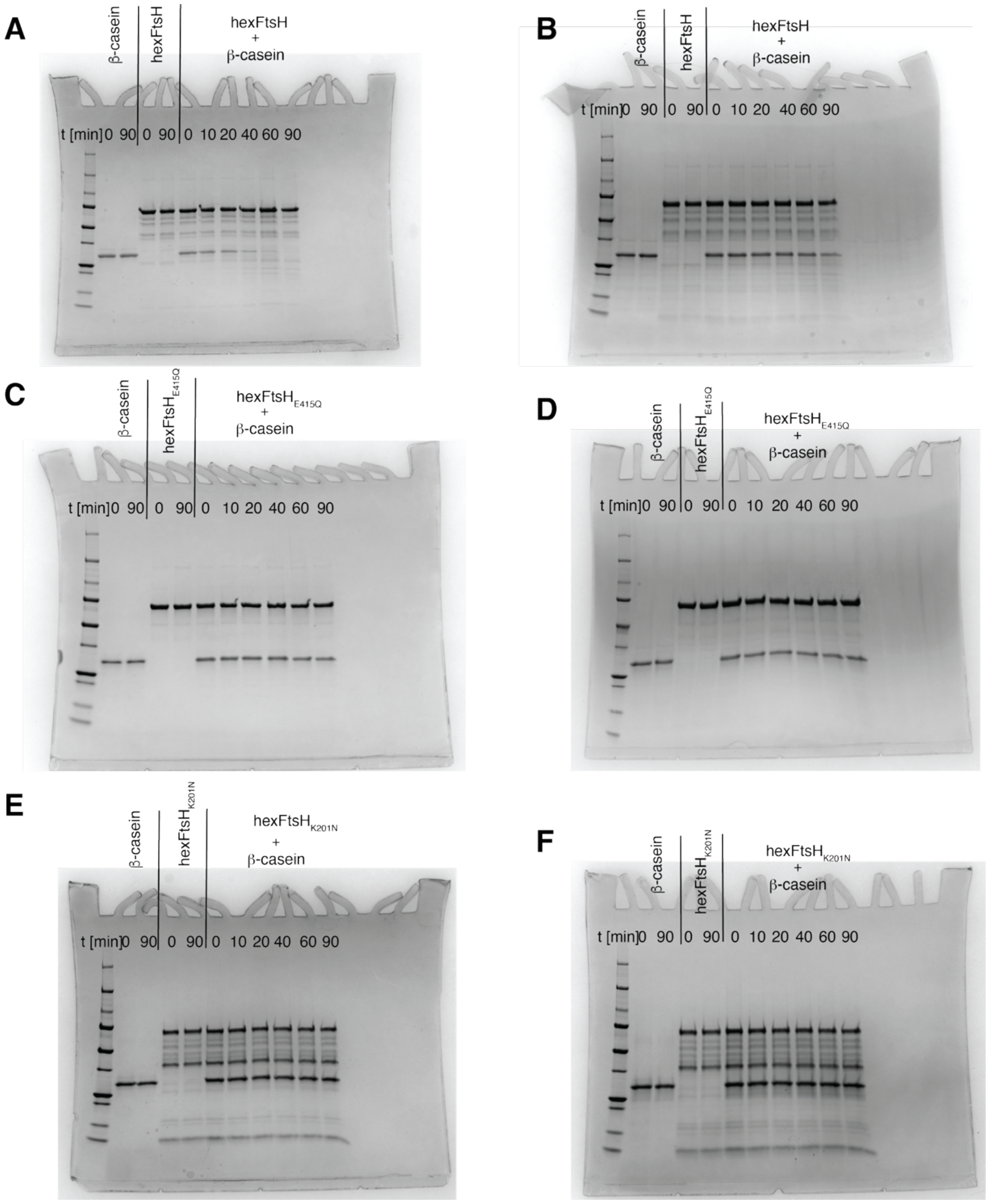


**Supplementary Figure S7.** Uncropped and unmodified gels underlying cleavage data presented in the main figures including control of β-casein at start and after 90 minutes (lane 1 and 2 after molecular weight ladder respectively) and control of FtsH variants at start and after 90 minutes (lane 3 and 4 respectively). **A)** hexFtsH + ATP. **B)** hexFtsH + ATPγS. **C)** hexFtsH$_{E415Q}$ + ATP. **D)** hexFtsH$_{E415Q}$ + ATPγS. **E)** hexFtsH$_{K201N}$ + ATP. **F)** hexFtsH$_{K201N}$ + ATPγS.

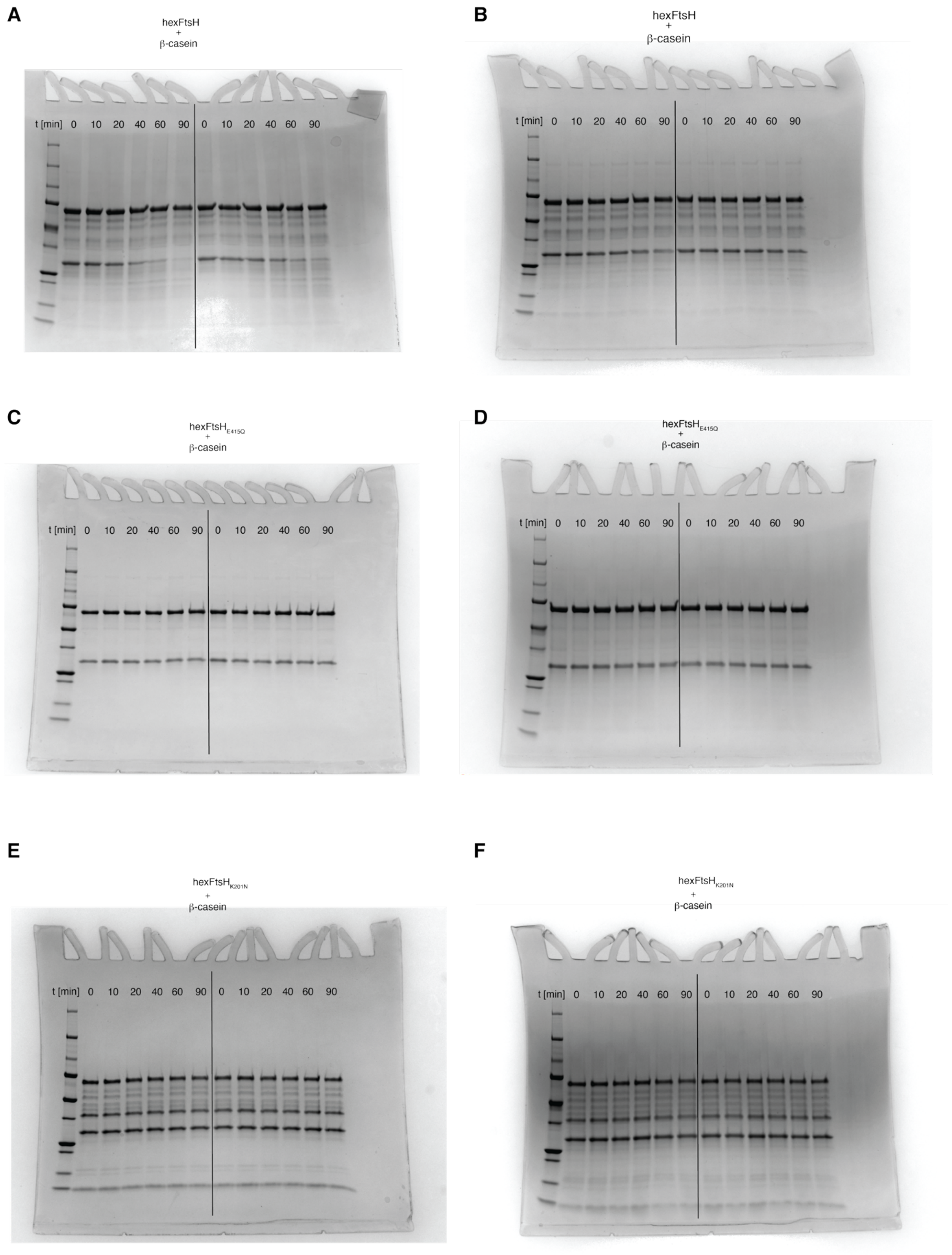


**Supplementary Figure S8.** Additional two replica experiments of the gels shown in **Supplementary Figure S7. A)** hexFtsH + ATP. **B)** hexFtsH + ATPγS. **C)** hexFtsH$_{E415Q}$ + ATP. **D)** hexFtsH$_{E415Q}$ + ATPγS. **E)** hexFtsH$_{K201N}$ + ATP. **F)** hexFtsH$_{K201N}$ + ATPγS.
